# A novel *Axin2* knock-in mouse model for visualization and lineage tracing of WNT/CTNNB1 responsive cells

**DOI:** 10.1101/2020.04.03.024182

**Authors:** Anoeska Agatha Alida van de Moosdijk, Yorick Bernardus Cornelis van de Grift, Saskia Madelon Ada de Man, Amber Lisanne Zeeman, Renée van Amerongen

**Affiliations:** Section of Molecular Cytology, Swammerdam Institute for Life Science, University of Amsterdam, Postbus 1212, 1000 BE, Amsterdam, the Netherlands

**Keywords:** beta-catenin, Axin2, doxycycline-inducible lineage tracing, GFP reporter mouse, organoids

## Abstract

Wnt signal transduction controls tissue morphogenesis, maintenance and regeneration in all multicellular animals. In mammals, the WNT/CTNNB1 (Wnt/β-catenin) pathway controls cell proliferation and cell fate decisions before and after birth. It plays a critical role at multiple stages of embryonic development, but also governs stem cell maintenance and homeostasis in adult tissues. However, it remains challenging to monitor endogenous WNT/CTNNB1 signaling dynamics *in vivo*. Here we report the generation and characterization of a new knock-in mouse strain that doubles as a fluorescent reporter and lineage tracing driver for WNT/CTNNB1 responsive cells. We introduced a multi-cistronic targeting cassette at the 3’ end of the universal WNT/CTNNB1 target gene *Axin2*. The resulting knock-in allele expresses a bright fluorescent reporter (3xNLS-SGFP2) and a doxycycline-inducible driver for lineage tracing (rtTA3). We show that the *Axin2*^*P2A-rtTA3-T2A-3xNLS-SGFP2*^ strain labels WNT/CTNNB1 cells at multiple anatomical sites during different stages of embryonic and postnatal development. It faithfully reports the subtle and dynamic changes in physiological WNT/CTNNB1 signaling activity that occur *in vivo*. We expect this mouse strain to be a useful resource for biologists who want to track and trace the location and developmental fate of WNT/CTNNB1 responsive stem cells in different contexts.

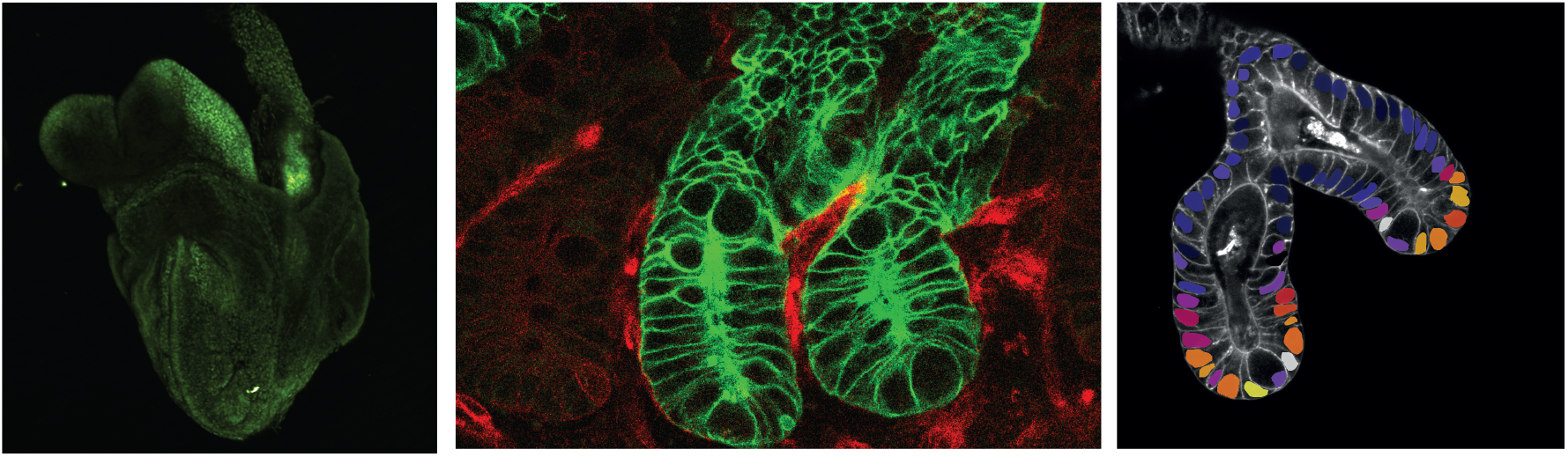

## Introduction

WNT/CTNNB1 (Wnt/β-catenin) signaling controls multiple stages of development in the mammalian embryo, ranging from primary axis formation (Liu et al., 1999) to mammary gland morphogenesis (Chu et al., 2004). Postnatally, WNT/CTNNB1 responsive stem cells regulate tissue homeostasis, regeneration and wound healing (Nusse & Clevers, 2017). Over the years, different genetically engineered mouse strains have been generated to capture these dynamic processes *in vivo* (Figure 1A).

**Figure 1.**
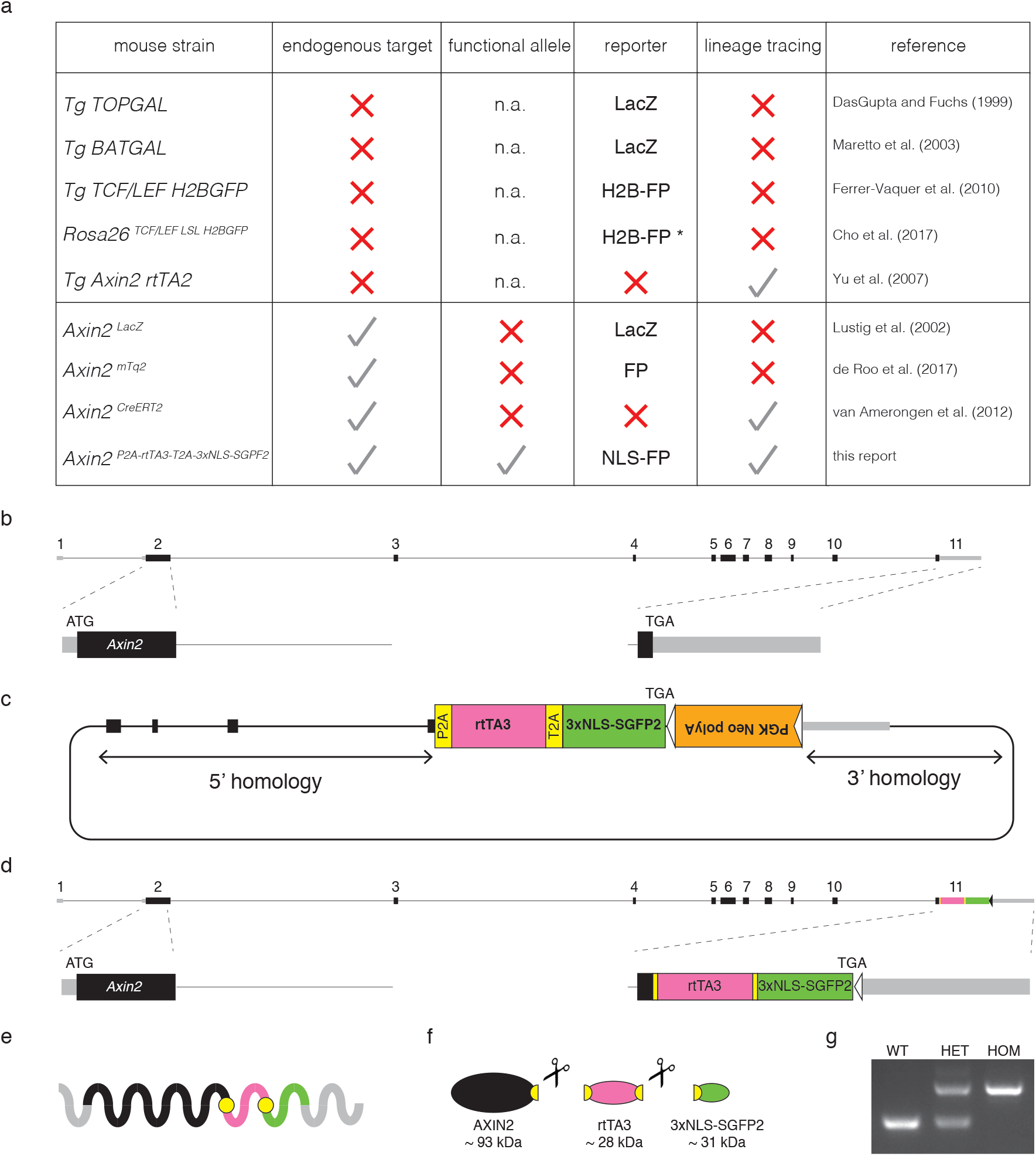
Design and generation of the *Axin2*^*P2A-rtTA3-T2A-3xNLS-SGFP2*^ knock-in allele. (A) Overview of WNT/CTNNB1 reporter and lineage tracing strains that are available from public repositories. Information was retrieved from the International Mouse Strain Resource (IMSR) at http://www.findmice.org. Strains are subdivided by four criteria: whether it is a random transgenic insertion (Tg) or a targeted insertion (Rosa26 or Axin2 locus); whether the targeted *Axin2* allele still produces a functional protein; whether the strain carries a reporter that can directly visualize WNT/CTNNB1 responsive cells; whether the strain can be used for lineage tracing experiments. Abbreviations: n.a.= not applicable, FP= fluorescent protein, *= requires Cre-mediated removal of a stop-cassette before the reporter becomes functional. (B) Schematic representation of the mouse *Axin2* locus (chr11:108920349-108950781, mm10 coordinates). Most existing *Axin2* knock-in strains target the 5’ end of the gene by introducing the knock-in cassette at the start codon (ATG) in exon 2. This disrupts the endogenous *Axin2* coding sequence. (C) Cartoon depicting the *Axin2*^*P2A-rtTA3-T2A-3xNLS-SGFP2*^ targeting construct. A multi-cistronic targeting cassette was cloned immediately upstream of the *Axin2* stop codon (TGA) in exon 11. The *PGK-Neo-polyA* cassette, flanked by *FRT* sites (indicated by triangles), was used for selection of embryonic stem cells and was removed prior to establishing the colony. (D) Targeted locus after removal of the *PGK-Neo-polyA* selection cassette. (E) The 3’ knock-in allele is designed to give rise to a single transcript in which the *Axin2* 5’ UTR, coding sequence and 3’ UTR are left intact. (F) Following translation, the self-cleaving P2A and T2A sequences ensure that the polypeptide is cleaved into a fully functional AXIN2 protein, a doxycycline activatable rtTA3 driver for lineage tracing, and a bright green fluorescent protein that localizes to the nucleus (3xNLS-SGFP2). (G) Genotyping PCR of wildtype, heterozygous and homozygous animals. Wildtype allele = 257 bp, knock-in allele = 461 bp.

Multiple reporter lines allow visualization of cells with active WNT/CTNNB1 signaling, either directly (e.g. GFP) or indirectly (e.g. lacZ). Some strains report active WNT/CTNNB1 signaling using an artificial reporter construct with concatemerized TCF/LEF sites, analogous to the first *in vivo* WNT/CTNNB1 reporter mouse, TOPGAL (Cho, Smallwood, & Nathans, 2017; DasGupta & Fuchs, 1999; Ferrer-Vaquer et al., 2010; Maretto et al., 2003). Although these transgenic strains continue to be highly useful, they likely do not fully recapitulate the endogenous pattern of WNT/CTNNB1 signaling activity due to integration of this reporter cassette into a random or exogenous locus (Cho et al., 2017; Yu, Liu, Costantini, & Hsu, 2007). Moreover, different expression patterns can be detected when multiple lines are compared in the same tissue (Al Alam et al., 2011). Although truly universal WNT/CTNNB1 target genes are rare, the negative feedback regulator *Axin2* has been shown to reliably report cells with active WNT/CTNNB1 signaling (Lustig et al., 2001). Accordingly, it can be used to track the developmental fate of WNT/CTNNB1 responsive stem cells in postnatal tissues (Van Amerongen, Bowman, & Nusse, 2012).

All existing models suffer from some drawbacks. Specifically, none of the available strains combine the expression of a sensitive fluorescent reporter gene and a lineage tracing driver from an unperturbed, physiological locus. Therefore, we generated a novel mouse strain, *Axin2*^*P2A-rtTA3-T2A-3xNLS-SGFP2*^, in which the knock-in cassette allows both direct visualization and doxycycline-dependent lineage tracing of WNT/CTNNB1 responsive cells. Our model faithfully recapitulates the endogenous *Axin2* expression pattern, while leaving normal *Axin2* expression intact.

## Results and Discussion

We designed our new *Axin2* knock-in allele to meet specific criteria. First, we targeted the 3’ end of the gene to maintain endogenous *Axin2* expression and to preserve both 5’ and 3’ regulatory control (Figure 1B-D, Supplementary Figure 1). Second, we wanted a single allele to function as both a reporter and a lineage tracing driver (Figure 1E-F). Third, we designed the reporter to serve as a direct readout of *Axin2* activity and to be suitable for live cell imaging. For this reason, we incorporated a fluorescent protein rather than a *lacZ* reporter gene. Based on prior experience, we expected *Axin2* expression to be relatively low. Therefore, we concentrated the reporter signal in the nucleus by fusing the *SGFP2* gene to a strong nuclear localization signal (3xNLS). Finally, for lineage tracing purposes we chose a doxycycline rather than a tamoxifen inducible system, because tamoxifen can affect the outgrowth of hormone responsive tissues such as the mammary gland (Shehata, van Amerongen, Zeeman, Giraddi, & Stingl, 2014), as well as impair embryonic development, pregnancy and delivery (Ved, Curran, Ashcroft, & Sparrow, 2019). To promote equimolar expression of AXIN2, rtTA3 and 3xNLS-SGFP2, self-cleaving 2A peptides were favored over IRES sequences (Goedhart et al., 2011).

Wildtype, heterozygous and homozygous *Axin2*^*P2A-rtTA3-T2A-3xNLS-SGFP2*^ mice were born at the expected mendelian ratios from heterozygous intercrosses on a C57BL/6 background (Figure 1G, Supplementary Table 1). We have not detected any phenotype (including differences in weight, fertility and lifespan up to 12 months) associated with homozygosity of the knock-in allele (data not shown). This is in contrast to 5’ *Axin2* knock-in alleles, which disrupt endogenous *Axin2* expression (de Roo et al., 2017; Lustig et al., 2001; Van Amerongen, Bowman, et al., 2012) and which can be hypomorphic on certain backgrounds (Supplementary Table 2). This results in skeletal phenotypes such as craniosynostosis (Yu et al., 2005) and typically prevents their use as homozygous reporter.

For functional validation of the allele, we isolated primary mouse embryonic fibroblasts (MEFs) from E13.5 embryos that were heterozygous for the *Axin2*^*P2A-rtTA3-T2A-3xNLS-SGFP2*^ allele. Only cells with activated WNT/CTNNB1 signaling are predicted to induce *Axin2* and to express rtTA3, as well as a bright green fluorescent protein in the nucleus (Figure 2A-B). Quantitative RT-PCR analysis confirmed that *Axin2*, *rtTA3* and *SGFP2* were all induced upon WNT/CTNNB1 pathway activation (Figure 2C-F). As expected, nuclear SGFP2 expression was induced upon dose-dependent activation of WNT/CTNNB1 signaling, as demonstrated by exposing the cells to increasing concentrations of CHIR99021 (a small molecule GSK3 inhibitor that activates WNT/CTNNB1 signaling downstream of the WNT receptor complex) (Figure 2G-H, Supplementary Figure 2).

**Figure 2.**
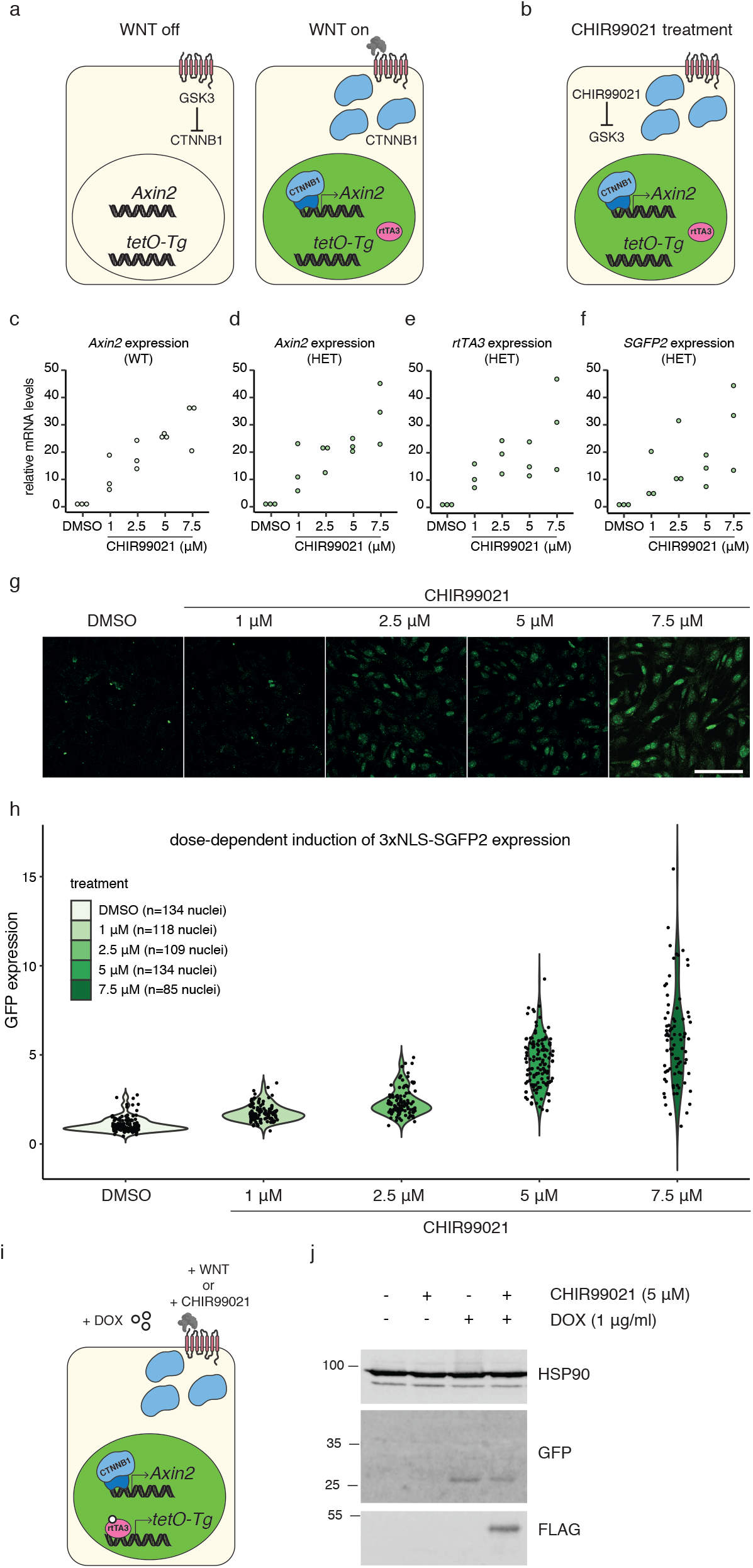
Functional validation of the *Axin2*^*P2A-rtTA3-T2A-3xNLS-SGFP2*^ allele. (A-B) Cartoons depicting the response of the knock-in allele to downstream WNT/CTNNB1 signaling, either by activation by WNT proteins *in vivo* (A) or by treatment with CHIR99021, a specific GSK3 inhibitor (B). Cells with active WNT/CTNNB1 signaling will induce Axin2, resulting in the expression of 3xNLS-sGFP expression (depicted as green nuclei). (C-F) Dot plots showing the dose-dependent induction of *Axin2* mRNA expression in wildtype (C) and heterozygous (D) mouse embryonic fibroblasts (MEFs), as measured by qRT-PCR in n=3 independent MEF isolates. *Rpl13a* was used as a reference gene, values in the DMSO treated control were set to 1. (E-F) Same as for (C-D), but showing *rtTA3* (E) and *SGFP2* (F) mRNA expression in heterozygous MEFs. (G) Confocal microscopy images of fixed mouse embryonic fibroblasts (MEFs), showing the dose-dependent induction and direct detection of SGFP2. Scalebar is 100 *μ*m. (H) Quantification of the experiment described and depicted in (G). Fixed MEFs were counterstained with DAPI to allow nuclear segmentation, after which the relative SGFP2 expression levels were calculated by correcting for the fluorescence intensity of the DAPI signal. This experiment was performed for n=3 independent MEF isolates. One experiment is shown here. The results for two additional MEF isolates are shown in Supplementary Figure 2. (I) The rtTA3 transcriptional activator is induced together with SGFP2, but only becomes active in the presence of doxycycline (DOX). This results in activation of *tetO*-driven transgenes (*tetO-Tg*). (J) Western blot showing the WNT/CTNNB1-dependent induction of SGFP2 and the WNT/CTNNB1- and DOX-dependent induction of a FLAG-tagged protein in whole cell lysates from MEFs carrying both the *Axin2* knock-in allele and a *tetO*-responsive transgene after treatment with different combinations of CHIR99021 and DOX.

Next, we tested if rtTA3 was capable of activating a *tetO*-responsive promoter in a doxycycline and CTNNB1-dependent manner (Figure 2I-J). In MEFs heterozygous for both the *Axin2*^*P2A-rtTA3-T2A-3xNLS-SGFP2*^ allele and a *tetO*-responsive *FLAG*-tagged transgene, FLAG-tagged protein expression was only induced in the presence of both CHIR99021 and doxycycline, but not by either treatment alone (Figure 2J).

To determine if the knock-in allele properly reports WNT/CTNNB1 signaling during embryonic development, we isolated embryos from stages at which defined *Axin2* expression domains were previously determined using RNA *in situ* hybridization (Jho et al., 2002). A dim SGFP2 signal could be detected in freshly isolated heterozygous or homozygous, but not wildtype embryos under a dissecting microscope (Supplementary Figure 3). Homozygous embryos consistently showed a brighter GFP signal than heterozygous embryos, suggesting bi-allelic expression of *Axin2*.

At E8.5, SGFP2 was expressed in the head folds and the posterior neural tube (Figure 3A-C). At E10.5, we detected SGFP2 expression in the developing limb bud, in the branchial arches, and along the dorsal neural tube, including the roof plate of the developing brain (Figure 3D-F). These are the same sites where endogenous *Axin2* was previously shown to be expressed via RNA *in situ* hybridization (Jho et al., 2002). At E12.5, prominent SGFP2 expression was visible in the mammary placodes (Figure 3G-I), recapitulating expression of the *Axin2*^*lacZ*^ reporter at this site (Van Amerongen, Bowman, et al., 2012). Using wholemount confocal microscopy, individual SGFP2-positive nuclei could readily be distinguished in these embryos at the indicated sites (Figure 3C, E, F, I, Supplementary Movie 1-2). Together, these experiments confirm that *Axin2*^*P2A-rtTA3-T2A-3xNLS-SGFP2*^ reports endogenous WNT/CTNNB1 signaling in the developing embryo.

**Figure 3.**
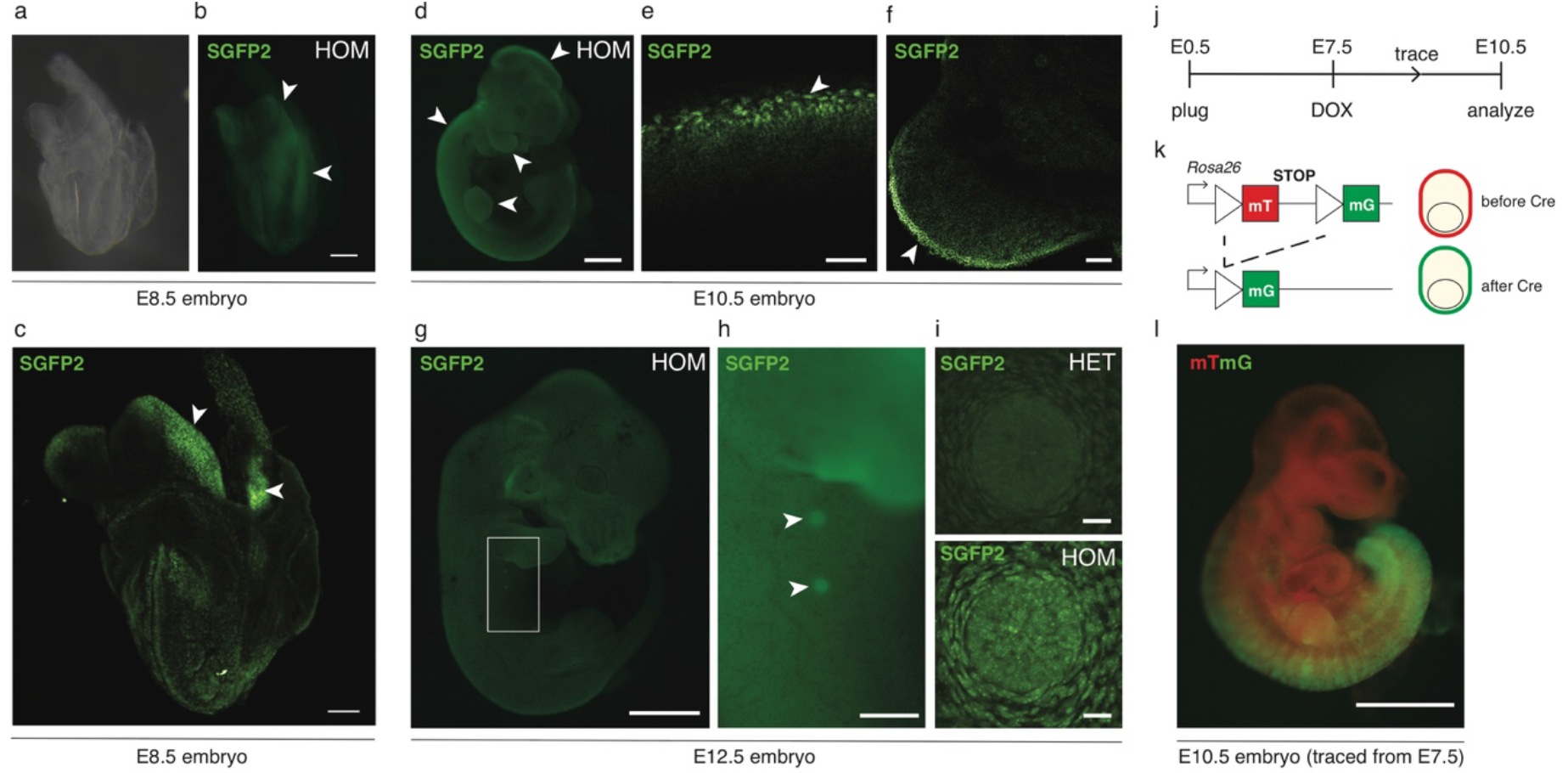
*Axin2*^*P2A-rtTA3-T2A-3xNLS-SGFP2*^ reports embryonic WNT/CTNNB1 signaling. (A-B) Wholemount E8.5 embryo after fixation, imaged under a dissecting microscope using brightfield (A) or fluorescent (B) settings. Arrowheads indicate the headfolds and posterior neural tube. Scalebar is 100 *μ*m. (C) Maximum intensity projection of a wholemount confocal microscopy Z-stack of the same embryo as in (A-B). Arrowheads point to regions with individual SGFP2-positive nuclei in the headfolds and posterior neural tube. Scalebar is 100 *μ*m. (D) Wholemount E10.5 homozygous embryo imaged under a fluorescence dissecting microscope. Arrowheads indicate expression in the developing limb bud and branchial arches, along the dorsal neural tube, and in the roof plate of the developing brain. Scalebar is 500 *μ*m. The same embryo is depicted in Supplementary Figure 2. (E-F) Wholemount confocal microscopy images of the roof plate (E) and limb bud (F) of an E10.5 heterozygous embryo. Individual SGFP2-positive nuclei can be distinguished and are indicated with arrowheads. Scalebar is 50 *μ*m for (E) and 100 *μ*m for (F). (G) Wholemount E12.5 homozygous embryo imaged under a fluorescence dissecting microscope. Scalebar is 1 mm. (H) Close up of the area highlighted in (G). Arrowheads indicate expression of the fluorescent reporter in the mammary buds. Scalebar is 500 *μ*m. (I) Maximum intensity projection of a wholemount confocal microscopy Z-stack of the E12.5 mammary bud, showing the difference in GFP intensity between a heterozygous (HET, top) and homozygous (HOM, bottom) embryo. Nuclear SGFP2 signal can be detected in both the epithelial bud and the surrounding mesenchyme. A 3D rotation of the entire Z-stack is provided in Supplementary Movies 1 (HET) and 2 (HOM). (J) Timeline for the experiment depicted in (L). Lineage tracing in WNT/CTNNB1-responsive cells was induced *in utero* at E7.5 by administering a single intraperitoneal injection of doxycycline (DOX) to pregnant females. Embryos were isolated and imaged at E10.5. (K) Lineage tracing principle using the *Rosa26*^*mTmG*^ reporter. Prior to Cre-mediated recombination, all cells express a membrane-localized red fluorescent protein (mT). Cre recombination results in a permanent switch to expression of a membrane-localized green fluorescent protein (mG). (L) Composite image showing *Cre/lox* dependent recombination in the posterior half of an *Axin2*^*P2A-rtTA3-T2A-3xNLS-SGFP2*^*;tetO-Cre;Rosa26*^*mTmG*^ triple heterozygous embryo. Red signal reflects non-recombined cells (mT), green signal reflects recombined cells (mG). Note that expression of membrane-bound GFP in the *Rosa26*^*mTmG*^ reporter is driven by a strong CAGGS promoter, such that with the imaging and filter settings chosen the weaker endogenous 3xNLS-SGFP2 signal of the *Axin2* knock-in allele itself is not detected. The embryonic timed matings and tracing experiments were performed in two independent litters for all timepoints. Scalebar is 500 *μ*m.

To test functionality of the lineage tracing module *in vivo*, we generated compound *Axin2*^*P2A-rtTA3-T2A-3xNLS-SGFP2*^;*tetO-Cre;Rosa26*^*mTmG*^ mice and labelled WNT/CTNNB1-responsive cells in E7.5 embryos by a single intraperitoneal injection of doxycycline into timed pregnant females. At E10.5, triple-heterozygous embryos showed prominent recombination of the *Rosa26*^*mTmG*^ reporter in the posterior half of the embryo (Figure 3J-K), consistent with posterior expression of *Axin2* at the time of induction at E7.5 (Jho et al., 2002). Thus, *Axin2*^*P2A-rtTA3-T2A-3xNLS-SGFP2*^ allows direct visualization and efficient lineage tracing of WNT/CTNNB1 responsive cells in the developing embryo.

Postnatally, SGFP2 expression was readily visualized in the small intestinal crypt. The strongest signal was found at the crypt bottom in so-called crypt-base-columnar cells (CBCs), which are known to be WNT/CTNNB1 responsive (Barker et al., 2007). These cells are readily identified by their distinct shape and presence among SGFP2-negative Paneth cells (Figure 4A-B). In adult mice, a gradient of SGFP2 signal extended into the upper portion of the crypt (Figure 4A). *Axin2* expressing cells in the intestine could also be detected by immunohistochemical staining of formalin fixed paraffin embedded (FFPE) tissue sections with an anti-GFP antibody (Supplementary Figure 4).

**Figure 4.**
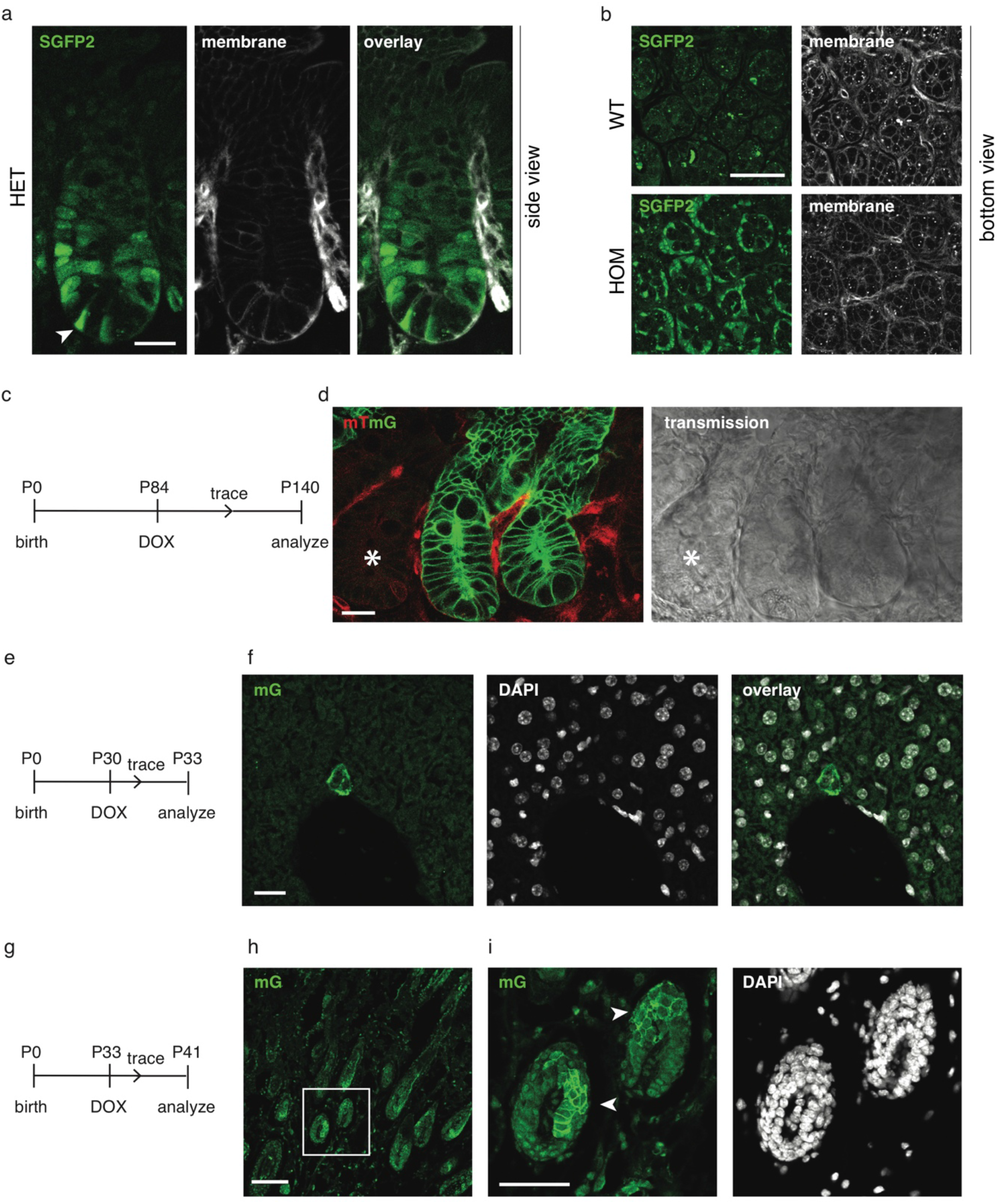
Visualization and lineage tracing of WNT/CTNNB1 responsive cells in postnatal tissues. (A) Wholemount confocal microscopy images, showing a side view of the intestinal crypts of adult heterozygous *Axin2*^*P2A-rtTA3-T2A-3xNLS-SGFP2*^ mice. Arrowhead indicates an SGFP2-positive stem cell, flanked by two SGFP2-negative Paneth cells. Scalebar is 20 *μ*m. (B) Wholemount confocal microscopy images, showing a bottom view of the intestinal crypts of wild type (WT, top) and homozygous (HOM) *Axin2*^*P2A-rtTA3-T2A-3xNLS-SGFP2*^ mice. In mice carrying the *Axin2* knock-in allele, direct SGFP2 expression (left) can be detected in stem cells at the bottom of the crypt, but not in the neighboring Paneth cells. Scalebar is 50 *μ*m. Cell outlines in (A) and (B) are visualized because the animals also express a membrane-bound tdTomato protein. (C) Timeline for the experiment depicted in (D). Lineage tracing in WNT/CTNNB1-responsive cells was induced in adult mice at P84 via a single intraperitoneal injection of doxycycline (DOX). The WNT/CTNNB1-responsive lineage was traced for 54 days. (D) Wholemount confocal microscopy images, showing a side view of the intestinal epithelium from an *Axin2*^*P2A-rtTA3-T2A-3xNLS-SGFP2*^*;tetO-Cre;Rosa26*^*mTmG*^ triple-heterozygous animal. Left panel shows the fluorescence signal coming from the *Rosa26*^*mTmG*^ lineage tracing reporter allele, with the membrane restricted red color (mT) reflecting non-recombined cells and the membrane restricted green color (mG) reflecting recombined cells. Right panel shows the transmission. Note that expression of membrane-bound GFP in the *Rosa26*^*mTmG*^ reporter is driven by a strong CAGGS promoter, such that with the imaging settings chosen the weaker endogenous 3xNLS-SGFP2 signal of the *Axin2* knock-in allele itself is not detected. Asterisk indicates a non-recombined crypt neighboring two traced crypts. Scalebar is 20 *μ*m. (E) Timeline for the experiment depicted in (F). Lineage tracing in WNT/CTNNB1-responsive cells was induced in pubertal mice at P30 via a single intraperitoneal injection of doxycycline (DOX). The WNT/CTNNB1-responsive lineage was traced for 3 days. (F) Confocal microscopy images of FFPE liver sections, showing the immunofluorescent detection of the recombined mTmG allele (left) in sporadic cells adjacent to the central vein. Nuclei are counterstained with DAPI (middle). Scale bar is 20 *μ*m. (G) Timeline for the experiment depicted in (H-I). Lineage tracing in WNT/CTNNB1-responsive cells was induced in pubertal mice at P33 via a single intraperitoneal injection of doxycycline (DOX). The WNT/CTNNB1-responsive lineage was traced for 8 days. (H) Confocal microscopy images of FFPE skin section, showing the immunofluorescent detection of the recombined mTmG allele in multiple hair follicles. Scale bar is 100 *μ*m. (I) Close up of the boxed area in (H), showing patches of recombined cells expressing membrane-localized mG (arrowhead) in the hair follicle (green, left). Nuclei are counterstained with DAPI (grey, right). Scale bar is 50 *μ*m. In these qualitative lineage tracing experiments, no difference was observed between male and female mice.

In adult triple-heterozygous *Axin2*^*P2A-rtTA3-T2A-3xNLS-SGFP2*^*;tetO-Cre;Rosa26*^*mTmG*^ mice, Cre/lox mediated recombination of the *Rosa26*^*mTmG*^ reporter allele in fast dividing, long-lived WNT/CTNNB1 responsive intestinal stem cells was efficiently induced by a single intraperitoneal injection of doxycycline in both pubertal and adult mice. Recombined cells first became visible 24-48 hours after doxycycline injection (Supplementary Figure 5), after which the progeny of fast dividing, long-lived WNT/CTNNB1-responsive stem cells could be traced and seen to populate the crypt and villus compartments within 3-6 days (Supplementary Figure 6). Prolonged tracing of the WNT/CTNNB1 responsive cell lineage resulted in sustained labeling of the entire crypt villus compartment as expected (Figure 4C-D), confirming that *Axin2*^*P2A-rtTA3-T2A-3xNLS-SGFP2*^ labels intestinal stem cells (Van Amerongen, Bowman, et al., 2012).

We and others have previously reported the presence of *Axin2*-positive, WNT/CTNNB1 responsive cells in multiple endodermal and ectodermal tissues (Lim et al., 2013; Lim, Tan, Yu, Lim, & Nusse, 2016; Van Amerongen, Bowman, et al., 2012; Wang, Zhao, Fish, Logan, & Nusse, 2015). Contrary to what is observed in the intestine, tracing of cells in other tissues was less efficient in the experimental set up used. Nevertheless, scarce labeling of cells in the liver (Figure 4E-F, Supplementary Movie 3) and prominent labeling of cells in the hair follicle (Figure 4G-I, Supplementary Figure 7) could be detected, either by imaging wholemount tissues or FFPE sections.

Finally, we established primary 3D small intestinal organoid cultures from *Axin2*^*P2A-rtTA3-T2A-3xNLS-SGFP2*^ animals. One heterozygous and one homozygous organoid line were maintained for extended cultures of more than 6 months, during which the reporter remained stably expressed and did not get silenced (Figure 5, Supplementary Figure 8). Similar to what we observed in embryos, the levels of SGFP2 were higher in homozygous than in heterozygous organoids (Supplementary Figure 8A-D). Strongest expression of the reporter allele was observed in CBCs at the bottom of the crypt-like compartment. Expression gradually decreased along the crypt axis in the transit amplifying compartment, similar to what is observed *in vivo* (Figure 4A, Figure 5A-D).

**Figure 5.**
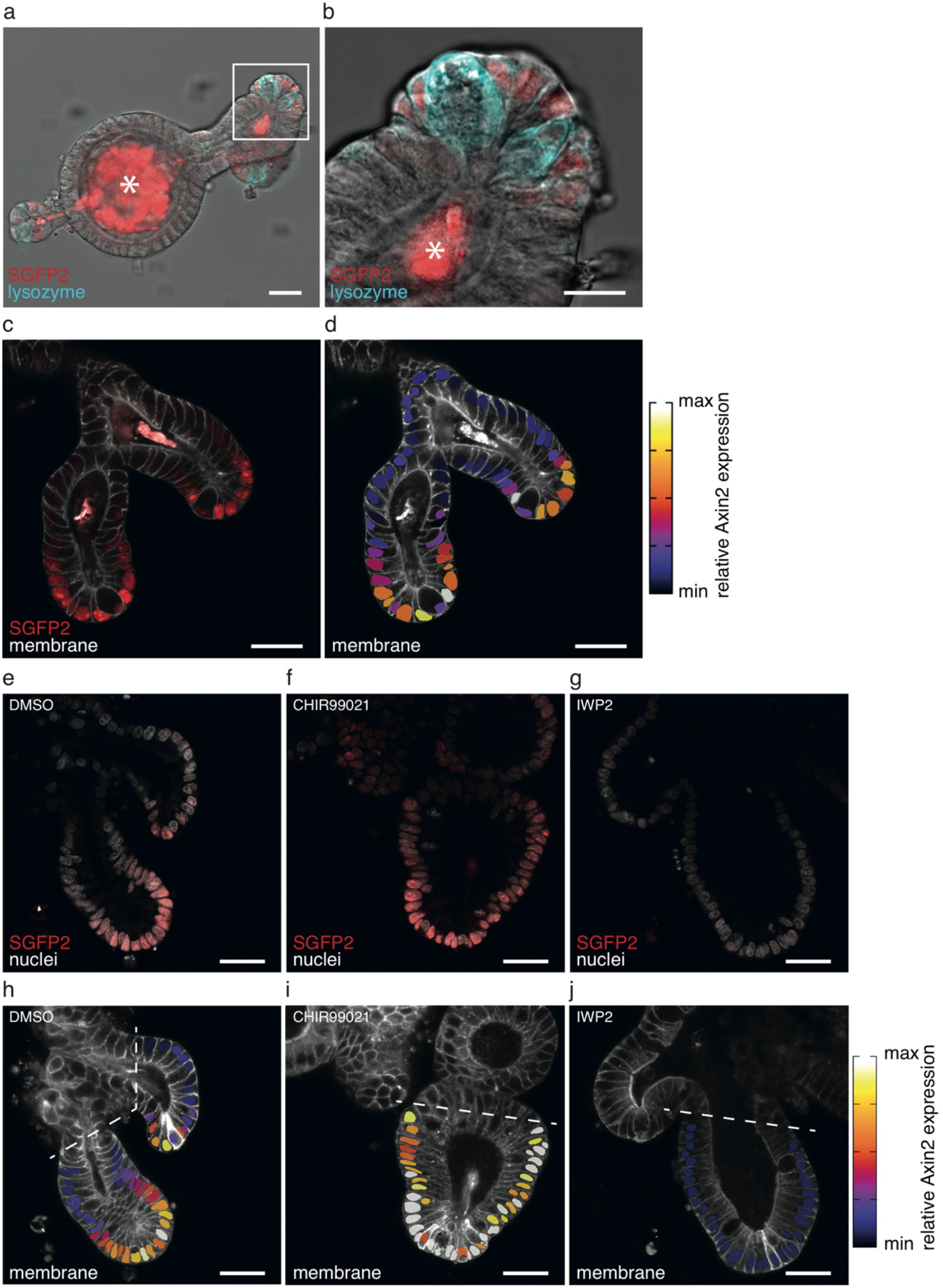
Visualization and manipulation of endogenous WNT signaling in small intestinal organoids. (A-B) Confocal microscopy (transmission) image of a fixed *Axin2*^*P2A-rtTA3-T2A-3xNLS-SGFP2 HOM*^*;Rosa26*^*mTmG HET*^ (red) small intestinal organoid stained with anti-Lysozyme (cyan). SGFP2 expression (depicted in red) is restricted to the crypt bottom, but does not co-localize with lysozyme staining in the large granular Paneth cells. The asterisk indicates dead material excreted into the organoid lumen that gives autofluorescence in the SGP2 channel. SGFP2 is depicted in a red look up table for better visual representation of the subtle differences in combination with nuclear and membrane stainings. Note that this organoid line is also heterozygous for the *Rosa26*^*mTmG*^ allele, which is used as a membrane marker in panels (C-J). (B) Close-up of the highlighted crypt section from (A). Scalebars are 30 *μ*m (A) and 15 *μ*m (B). For (A-B) a total of 17 organoids were imaged in a single experiment. A representative image is shown. (C) Confocal microscopy image of two crypts from a fixed *Axin2*^*P2A-rtTA3-T2A-3xNLS-SGFP2 HOM*^*;Rosa26*^*mTmG HET*^ small intestinal organoid. Membrane-tdTomato is depicted in grey and 3xNLS-SGFP2 expression in red. 3xNLS-SGFP2 is expressed in a gradient along the crypt-axis, the highest expression is located at the crypt bottom and gradually declines when cells move upward. Scalebar is 30 *μ*m. (D) Heat map showing the relative *Axin2* expression superimposed on the confocal image in Figure 5C. The heat map depicts the 3xNLS-sGPF2 signal relative to a DRAQ5 nuclear staining (not shown) and is imposed on the area of the nuclei. This overlay corrects for signal intensity differences due to imaging depth and reveals differences in gene expression. Scalebar is 30 *μ*m. For panels (C-D) a total of 14 organoids were imaged in 2 independent experiments. A representative image is shown. (E-F) Confocal microscopy images of crypts from fixed *Axin2*^*P2A-rtTA3-T2A-3xNLS-SGFP2 HOM*^*;Rosa26*^*mTmG HET*^ (3xNLS-SGFP2, red) small intestinal organoids stained with nuclear dye DRAQ5 (grey). The organoids were treated with either DMSO for 24 hours (E), CHIR99021 for 24 hours (F), or IWP2 for 48 hours (G) before fixation. Scalebar is 30 *μ*m. (H-J) Heat map showing relative *Axin2* reporter levels superimposed on the confocal images of (E-F). The dotted lines indicate the cutoff for which the nuclei were excluded from further analysis. Note that this relative scale is not directly comparable to the one depicted in (D). Scalebar is 30 *μ*m. For panels (E-J) a total of 51 organoids were imaged in 2 independent experiments (n=14 for DMSO, n=29 for CHIR99021, n=8 for IWP2). Representative images of all conditions are shown.

Intestinal organoids provide a relevant physiological context to study the bidirectional response of the reporter, since WNT/CTNNB1 signaling can be both hyperactivated and inhibited in this setting. Direct modulation of the activity of the WNT/CTNNB1 pathway with small molecule inhibitors influenced both the intensity and the number of SGFP2-positive cells present per organoid (Figure 5E-J). Exposure to CHIR99021 resulted in a prominent increase in *Axin2*-positive cells along the crypt axis within 24 hours (Figure 5F, I). In contrast, incubation with the PORCN inhibitor IWP-2, which blocks the secretion of endogenous WNT proteins, completely abolished SGFP2 expression within 48 hours (Figure 5G, J). This confirms that the *Axin2*^*P2A-rtTA3-T2A-3xNLS-SGFP2*^ reporter is sensitive to direct changes in WNT/CTNNB1 signaling and dynamically reports endogenous WNT/CTNNB1 signaling in individual cells.

In conclusion, our *Axin2*^*P2A-rtTA3-T2A-3xNLS-SGFP2*^ model offers multiple advantages over existing mouse models (Figure 1A) and as such we expect it to be useful for investigations into the role of WNT/CTNNB1 signaling in embryonic development and adult tissue stem cells. We show that the fluorescent 3xNLS-SGFP2 reporter is expressed at multiple embryonic and postnatal stages of development (Figure 3, 4) and dynamically reports changes in WNT/CTNNB1 signaling (Figure 2, 5). Signal intensity of the reporter can be increased by using homozygous rather than heterozygous mice.

We also demonstrate that the new, doxycycline-inducible lineage tracing driver is suitable for embryonic and postnatal traces (Figure 3, 4). At the same time, our model does not fully recapitulate the results we previously obtained with our 5’ *Axin2*^*CreERT2*^ knock-in model (Lim et al., 2013; Van Amerongen, Bowman, et al., 2012; Wang et al., 2015). This once again underscores that care should be taken when interpreting results from genetic reporters and, in general, when comparing expression patterns across different mouse strains. The discrepancy between the different *Axin2* lineage tracing models can be explained by one or more of the following reasons. First, our 3’ knock-in fully recapitulates endogenous *Axin2* expression and thereby reveals how low these expression levels really are, especially after birth. This is in agreement with the low-level and low-amplitude oscillations of *Axin2* expression detected by others (Sonnen et al., 2018) and something that may have been obscured by the enzymatic amplification that occurs during X-gal staining of *Axin2*^*LacZ*^ mice. Second, the *tetO-Cre* line used for our studies is not leaky, but may display a high threshold to activation. In combination with the low *Axin2* (and thus low *rtTA3*) expression this may result in insufficient Cre activity to recombine the *Rosa26*^*mTmG*^ allele. Whether *rtTA3* or *tetO-Cre*, or a combination of both, is the rate limiting factor remains to be tested. Labeling efficiency might be increased in homozygous *Axin2*^*P2A-rtTA3-T2A-3xNLS-SGFP2*^ animals or in combination with a different *tetO-Cre* strain.

Summarizing, our bright fluorescent nuclear SGFP2 reporter offers straightforward detection of *Axin2* expression and is expected to be compatible with live cell imaging, nuclear segmentation and thus automated cell tracking in embryonic tissues and organoids. The *Axin2*^*P2A-rtTA3-T2A-3xNLS-SGFP2*^ reporter is sensitive to direct changes in endogenous WNT/CTNNB1 signaling and allows monitoring of WNT/CTNNB1 sensitive cells. As such, our model can be a useful tool for the scientific community to simultaneously visualize and trace cells with active WNT/CTNNB1 signaling both *in vivo* and *in vitro*.

## Acknowledgements

We thank the van Leeuwenhoek Centre for Advanced Microscopy (LCAM, Section Molecular Cytology, Swammerdam Institute for Life Sciences, University of Amsterdam) for the use of their facilities and LCAM staff for sharing their expertise and providing technical support, Jeroen van Zon (AMOLF) for sharing the anti-Lysozyme antibody, all colleagues for stimulating discussions and Tanne van der Wal, Larissa Mourao and Dorus Gadella for feedback on the manuscript.

This work was supported by a MacGillavry fellowship from the University of Amsterdam (to RvA), a career development award from KWF Kankerbestrijding (2013-6057, to RvA), a research grant from KWF Kankerbestrijding/Alpe d’HuZes (2015-8014, to RvA) and an NWO-ALW VIDI grant (864.13.002, to RvA).

## Methods

### Generation of *Axin2*^P2A-rtTA3-T2A-3xNLS-SGFP2^ mice

The following considerations were made in designing the *Axin2*^*P2A-rtTA3-T2A-3xNLS-SGFP2*^ targeting construct: P2A and T2A self-cleaving peptides were selected to ensure optimal separation of Axin2, rtTA3 and 3xNLS-SGFP2 (Daniels, Rossano, Macleod, & Ganetzky, 2014). The rtTA3 (Tet3G) transactivator was selected to ensure sensitive and tight (i.e. low background) doxycycline-dependent control for lineage tracing purposes (Das et al., 2004; Zhou, Vink, Klaver, Berkhout, & Das, 2006). The SGFP2 fluorescent protein was selected for its bright and monomeric properties (Kremers, Goedhart, Van Den Heuvel, Gerritsen, & Gadella, 2007). In anticipation of low expression levels, a triple nuclear localization signal (3xNLS) was employed to concentrate the fluorescent reporter signal for improved detection (Chertkova et al., 2017).

The *Axin2*^*P2A-rtTA3-T2A-3xNLS-SGFP2*^ mouse line was generated by Ozgene Pty Ltd (Bentley WA, Australia). The final targeting construct, containing 5.5 kb and 2.8 kb homology arms, was linearized and electroporated into Bruce 4 [B6.Cg-Thy1] ES cells (Köntgen, Süss, Stewart, Steinmetz, & Bluethmann, 1993). Homologous recombinant ES cell clones (7/95) were identified by Southern hybridization and 2 independent clones were injected into goGermline blastocysts (Koentgen et al., 2016). Male chimeric mice were obtained and crossed to FLP females (OzFlp; a Rosa26 transgenic FLP deleter strain) to establish heterozygous germline offspring on a pure C57BL/6 background. The final colony was established from one of these ES cell derived chimeras. Cloning, sequencing and targeting details are available via <u>shorturl.at/ilqIO</u>.

### Mice

All mice used for this study were maintained under standard housing conditions. Animals were housed in open or IVC cages on a 12h light/dark cycle and received food and water *ad libitum*. All experiments were performed in accordance with institutional and national guidelines and regulations and approved by the Animal Welfare Committee of the University of Amsterdam.

*Axin2*^*P2A-rtTA3-T2A-3xNLS-SGFP2*^ mice were backcrossed to C57BL/6JRccHsd (Envigo) or FVB/NHan®Hsd (Envigo). These mice are currently being deposited with EMMA/JAX (official strain nomenclature: B6(Cg)-Axin2<tm1Rva>/Rva MGI:6360059). Most experiments were performed on a mixed C57BL/6 and FVB background, unless noted otherwise.

Other strains used: *Rosa26*^*mTmG*^ (mixed background, official strain name B6.129(Cg)- Gt(ROSA)26Sortm4(ACTB-tdTomato,-EGFP)Luo/J, Jackson labs stock number 007676, (Muzumdar, Tasic, Miyamichi, Li, & Luo, 2007)); *tetO-Cre(Bjd)* (mixed background, official strain name Tg(tetO-cre)LC1Bjd, Infrafrontier EMMA ID 00753, (Schönig, Schwenk, Rajewsky, & Bujard, 2002)); *tetO-FLAG-Wnt5a* (FVB background, official strain name FVB/N-Tg(tetO-Wnt5a)17Rva/J, Jackson labs stock number 022938, (Van Amerongen, Fuerer, Mizutani, & Nusse, 2012)); *Axin2*^*CreERT2*^ (mixed background, official strain name B6.129(Cg)-Axin2tm1(cre/ERT2)Rnu/J, Jackson labs stock number 018867, (Van Amerongen, Bowman, et al., 2012).

For timed matings, female mice were screened for the presence of a vaginal plug. When a plug was found, this timepoint was recorded as E0.5. For embryo isolation, pregnant dams were euthanized at the indicated timepoints and embryos were isolated for further processing. Yolk sacs were used for genotyping all embryos.

For embryonic lineage tracing, pregnant mice were injected intraperitoneally with 375 *μ*g doxycycline (75 *μ*l of a filter sterilized, 5 mg/ml solution of Doxycycline Hyclate (Merck) dissolved in PBS). For postnatal lineage tracing, mice were injected intraperitoneally with a filter sterilized solution of 2 mg doxycycline (Merck), dissolved in PBS (100 *μ*l or 200 *μ*l of a filter sterilized, 20 mg/ml or 10 mg/ml stock solution). We verified that no leakiness (i.e. recombination of the *Rosa26*^*mTmG*^ reporter in the absence of doxycycline) was detected in uninjected triple heterozygous animals. As negative controls for imaging experiments, we used littermates lacking the *tetO-Cre* allele, and thus also not showing recombination of the *Rosa26*^*mTmG*^ reporter.

### PCR Genotyping

Ear clips or yolk sacs were lysed either overnight (ear clips) or for 2 hours (yolk sacs) at 55°C in 200 *μ*l of Direct PCR tail lysis buffer (Viagen) supplemented with 200 *μ*g/ml Proteinase K (20 mg/ml stock solution). Proteinase K was inactivated by incubating the samples at 85°C for 15-45 minutes. Samples were cooled to room temperature and spun down (2 minutes at 14,000 rpm), after which 1 *μ*l of the supernatant was used as input for a standard 20 *μ*l PCR reaction with 0.4 *μ*l of Phire II polymerase (ThermoFisher, #F-124S). PCR conditions were as follows: initial denaturation at 98°C for 30 seconds, followed by 30 or 35 cycles of denaturation at 98°C for 5 seconds, annealing at the relevant temperature for 5 seconds, extension at 72°C for 10 seconds, followed by a final extension step of 72°C for 1 minute. Samples were cooled to 16°C and analyzed on a 2% agarose gel in standard TAE buffer. Primer sequences, annealing temperatures, number of cycles and expected band sizes are detailed in Supplementary Table 3.

### Isolation and culture of mouse embryonic fibroblasts

Mouse embryo fibroblasts (MEFs) were isolated at E13.5 as described previously (Van Amerongen et al., 2005). MEFs were cultured in DMEM supplemented with Glutamax (ThermoFisher Scientific), 10% FBS (ThermoFisher Scientific), 1% penicillin/streptomycin (ThermoFisher Scientific) and 50*μ*M β-mercaptoethanol (Merck) at 37°C and 5% CO2. To induce WNT/CTNNB1 signaling in these cells, CHIR99021 (BioVision) was dissolved in DMSO (Sigma Aldrich) at 6 mM and added at the concentrations described.

### Isolation and culture of mouse intestinal organoids

Mouse intestinal organoids were established as previously described (Sato et al., 2009). Crypts were isolated from the entire length of the small intestine of one *Axin2*^*P2A-rtTA3-T2A-3xNLS-SGFP2(HOM)*^;*Rosa26*^*mTmG(HET)*^ animal (used for figure 5), one *Axin2*^*P2A-rtTA3-T2A-3xNLS-SGFP2(HOM)*^ animal and one *Axin2*^*P2A-rtTA3-T2A-3xNLS-SGFP2(HET)*^ animal (used for the comparison of heterozygous and homozygous organoids in supplementary figure 8) and used to establish individual organoid lines. Organoids were cultured in 10 *μ*l Matrigel droplets (Corning) in culture medium containing advanced DMEM/F12 (ThermoFisher Scientific) supplemented with 100 U/ml Penicillin/Streptomycin (ThermoFisher Scientific), 2 mM Glutamax (ThermoFisher Scientific), 10mM HEPES (ThermoFisher scientific), 1x B27 supplement (ThermoFisher Scientific) and 1.25 mM N-acetylCysteine (Sigma Aldrich), freshly added EGF (50 ng/ml, PeproTech), recombinant murine Noggin (100 ng/ml, PeproTech) and recombinant murine R-spondin 1 (500 ng/ml, Sinobiological Inc.) at 37°C and 5% CO2. For passaging, cell culture medium was removed and Matrigel was broken into small pieces by scraping, followed by vigorous pipetting with ice-cold advanced DMEM/F12. Crypts were centrifuged at 200g for 5 min at 4°C. The supernatant was carefully removed, the pellet was resuspended in Matrigel and plated on pre-warmed plates. Medium was refreshed every other day and organoids were split once a week in a 1:3 ratio. To modulate the levels of WNT/CTNNB1 signaling at day 5 after plating, 10 *μ*M CHIR99021 (BioVision), or 2 *μ*M IWP-2 (Calbiochem) were added for 24 or 48 hours before fixation.

### RNA isolation, cDNA synthesis and qRT-PCR analysis

Total RNA was isolated from confluent MEF cultures using TRIzol (ThermoFisher Scientific) according to the manufacturer’s guidelines. Residual genomic DNA was removed by RQ1 RNAse-free DNAse treatment (Promega) according to the manufacturer’s instructions. The RNA concentration was determined using a Nanodrop spectrophotometer. cDNA was synthesized from 2 *μ*g RNA using SuperScript IV Reverse Transcriptase (Invitrogen) and Random Hexamers (ThermoFisher Scientific), according to the manufacturer’s instructions. RiboLock RNAse-inhibitor (ThermoFisher Scientific) was added during the reverse transcriptase reaction. The resulting cDNA was diluted 10-fold for subsequent qRT-PCR analysis.

qRT-PCR was performed using a QuantStudio 3 (Applied Biosystems). PCR reactions (total volume 20 *μ*l) were set up containing 13 *μ*l RNAse-free H2O, 4 *μ*l 15x HOT FIREPol EvaGreen qRT-PCR Mix Plus ROX (Solis Biodyne), 0.5 *μ*l of each specific forward and reverse primer (10 *μ*M stock) and 2 *μ*l of diluted cDNA template. The reactions were set up in technical triplicates in 96-well qPCR plates. One negative control (no-RT) reaction was included for each sample/primer combination. Thermal cycling was performed, starting with a hold stage at 50°C for 2 min, an initial step at 95°C for 15 min, followed by 40 cycles of denaturation at 95°C for 15 s and annealing at 60°C for 1 min. Each run was completed with a melting curve analysis. Primer sequences are detailed in Supplementary Table 3.

### Protein isolation and Western blot analysis

Cells were harvested by lysis in RIPA buffer (150mM NaCl, 1% NP-40, 0.5% sodium deoxycholate, 0.1% SDS, 50mM Tris-HCl, pH 8.0) supplemented with protease inhibitors (Roche). Protein concentration was determined using a Pierce BCA protein kit (BioRad), following the manufacturer’s instructions. Samples were prepared in equal amounts of protein in loading buffer (125 mM Tris-HCl (pH 6.8), 50% glycerol, 4% SDS, 0.2% Orange-G, 10% betamercaptoethanol (Merck)) and boiled at 95°C. Denatured samples were run on a 10% SDS-PAGE gel and transferred to a 0.2 *μ*M nitrocellulose membrane (Biorad) at 30V for 16 hours; or at 260mA for 3 hours, both at 4°C. The blot was blocked for 1 hour at room temperature in Odyssey Blocking Buffer (LI-COR) diluted 1:1 in TBS, followed by overnight incubation with primary antibody at 4°C. Primary antibodies were used to recognize GFP (1:1000, Thermo Fisher, A-6455), HSP90α/β (1:1000, Santa Cruz Biotechnology, sc-13119), Flag M1 (1:2000, Sigma, F3040), and active CTNNB1 (1:1000, Cell Signaling Technology, 8814). After incubation with the primary antibody, blots were washed extensively with TBS-Tween (0.1%) before incubating with the secondary antibodies for 1 hour at room temperature. Secondary antibodies were IRDye 680LT anti-mouse (1:20000, LI-COR) and IRDye 800CW anti-rabbit (1:20000, LI-COR). For Flag M1 antibody, 1mM calcium was added to all solutions from the addition of the 1^st^ antibody on. Blots were imaged with an Odyssey Fc Dual-Model Imaging System (LI-COR). Image visualization/representation for Western blots was performed using LI-COR Image Studio Lite software. Panels were cropped in Photoshop. Original Western blots are available via <u>shorturl.at/ilqIO</u>.

### Immunofluorescence and Immunohistochemistry

GFP expression was assessed using standard immunostaining methods. 4% PFA-fixed (Merck) and paraffin-embedded tissues were cut into 15 *μ*m sections (IHC) or 5 *μ*m sections (IF) and floated on water. Tissues sections were picked up onto a superfrost plus slide (Thermo Scientific), deparaffinized, and then rehydrated. Antigen retrieval was performed in Sodium Citrate buffer (10mM Sodium Citrate, 0,05% Tween 20, pH 6.0) by incubating the slides for 2.5 hours at 85°C, followed by cooling on ice. The slides were then washed with PBS for 5 minutes.

For immunohistochemistry, the slides were rinsed with demi water. Endogenous peroxidase activity was blocked by incubation in 0.3% hydrogen peroxide in methanol for 30 minutes, and then the tissues sections were rinsed with PBS. Next, tissues were circled with an ImmEdge pen (Vectorlabs) and incubated for 20 minutes with diluted normal blocking serum from Vectastain Elite ABC kit (PK-6101 Vector labs). Diluted GFP antibody (rabbit, GFP polyclonal antibody, A-6455 Invitrogen), was prepared by using 1:5000 GFP antibody in blocking serum. Without rinsing the slides, this antibody was incubated for 30 minutes at room temperature. As a negative control, samples were only stained with the secondary antibody. The slides were washed with PBS for 5 minutes for 3 times. Then the secondary antibody was added for 30 minutes, followed by 3 washes with PBS. At this point, Vectastain ABC reagent was prepared according to manufacturer’s instructions, and allowed to stand for 30 minutes before use. The ABC reagent was added to the slides for 30 minutes, followed by 3 washes with PBS.

Slides were rinsed in demi water and DAB peroxidase (ImmPACT HRP SK-4105) was added for 2-3 minutes, followed by a demi water wash. Sections were counterstained with 50% hematoxyilin, dehydrated in a graded series of ethanol dilutions followed by Histoclear II (National Diagnostics), and mounted with a coverslip using Omnimount (National Diagnostics).

For immunofluorescence, the slides were rinsed with PBS and tissues were circled with an ImmEdge pen (Vectorlabs) and incubated for 45 minutes with 2.5% BSA in PBS. Without rinsing the slides, the GFP antibody (chicken, GFP polyclonal antibody Abcam ab13970), diluted 1:400, was incubated over night at room temperature. As a negative control, samples were only stained with the secondary antibody. The slides were washed with PBS for 5 minutes for 3 times. Then the secondary antibody, Alexa Fluor 488 (Invitrogen A11039), diluted 1:1000, and DAPI (Invitrogen D1306, used at 1 ug/ml, stock solution 5 mg/ml in dimethylformamide) was added for 1 hour, followed by 3 washes with PBS. Next the slides were mounted with a coverslip using Mowiol (6 g glycerol, 2.4 gr polyvinylalcohol 4-88 (Sigma, 81381), 6 ml MQ and 12 ml 0.2 M Tris HCL pH 8.5).

### Microscopy

For imaging MEFs in the presence of DAPI, cells were fixed for 15 minutes in fresh 4% PFA made from 16% PFA (methanol free ampules, Thermo Scientific) diluted in PBS, and mounted in Mowiol with 1 *μ*g/ml DAPI (Invitrogen). Fixation was carried out at 37°C to prevent quenching of the endogenous fluorescence signal. Fixed cells were imaged using a Nikon A1 microscope. Fluorophores were excited as follows: DAPI at 405nm, SGFP2 at 488nm. Emission was detected as follows: DAPI 425-480nm, SGFP2 500-555. Images were acquired using a 40x oil objective.

For wholemount confocal microscopy of embryos and tissues, samples were fixed in 4% formaldehyde for histology, buffered pH 6.9 (Merck) or 4% PFA made from 16% PFA (methanol free ampules, Thermo Scientific) diluted in PBS for 1 hour at room temperature. Tissues were dehydrated through a graded ethanol series and cleared in methylsalicylate (Sigma) as described previously (Van Amerongen, Bowman, et al., 2012) or through a graded glycerol series, as indicated in the figure legends. Imaging of whole-mount embryos, intestine, liver and skin was performed on a Leica SP8 confocal microscope. Fluorophores were excited as follows: DAPI at 405nm, SGFP2 at 488nm, tdTomato at 561nm. Emission was detected as follows: DAPI 425-480nm, SGFP2 500-555nm, tdTomato 567-730nm. All images were acquired using a 10x dry, 20x dry or 40x oil objective.

Images of intestinal organoids were captured using confocal microscopy on a Leica SP8 with the LasX software. For imaging, the samples were cultured and imaged on glass chamber slides (Ibidi). Intestinal organoids were fixed in 4% formaldehyde for histology, buffered pH 6.9 (Merck) for 15 minutes at room temperature, the reaction was quenched with 0.15 M Glycine and the samples were permeabilized with 0.5% TritonX-100 for 10 minutes. The organoids were either incubated with 1 *μ*M DRAQ5 for 10 minutes at room temperature, or blocked with 5% BSA blocking solution for 2 hours. To visualize Paneth cells the samples were stained with 1:500 primary antibody anti-lysozyme (Agilent) overnight at 4°C, and 1:1000 secondary antibody Alexa Fluor 647 (Invitrogen) for 1 hour at room temperature. Fluorophores were excited as follows: sGPF2 at 488nm, tdTomato at 561nm, Alexa Fluor 647/anti-lysozyme and DRAQ5 at 633nm. Emission was detected as follows: SGFP2 HyD 494-540nm, tdTomato HyD 568-625nm, Alexa Fluor 647/anti-lysozyme PMT 642-696nm, DRAQ5 PMT 665-740nm. All images were acquired using a 40x oil or 63x water objective.

For imaging immunohistochemistry slides, pictures were taken using an Axio scope A1 microscope with a Nikon Ri2 camera and NIS F freeware. For imaging immunofluorescence slides, Pictures were taken using a Nikon A1 confocal microscope and NIS elements AR software.

Wholemount images of embryos were taken on a Leica stereomicroscope MZFLIII equipped with a Nikon Digital sight DS-Fi2 and NIS elements F freeware software.

### Microscopy Image analysis

Fluorescence microscopy images were processed in Fiji (Schindelin et al., 2012) using the Image 5D plugin and custom built approaches. For the quantification of SGFP2 expression in MEFs (Figure 2C-D and Supplementary Figure 2), ROIs surrounding single nuclei were selected manually in the DAPI channel using the magic wand tool. The SGFP2 nuclear signal was normalized over DAPI intensity. For each condition, 3 images were analyzed. This was repeated for 3 independent MEF lines. Data for one of the lines is shown in Figure 2, data for the other two lines are shown in Supplementary Figure 2. For Figure 5, heatmaps were drawn as follows: ROIs surrounding single nuclei were selected manually in the DRAQ5 channel using the Freehand selection tool. The SGFP2 nuclear signal was normalized over DRAQ5 intensity. To determine the relative *Axin2* expression a ratio of SGFP2 over DRAQ5 was calculated in Excel and imported in Fiji. An overlay was made that draws a heat map in selected ROIs based on this ratio with the ROI Color Coder plugin (Tiago Ferreira et al. 2016).

Color scheme choices: SGFP2 signal is shown in green, except for Figure 5 and Supplementary Figure 6, where a red LUT was chosen to be able to overlay the SGFP2 and the nuclear signal. All lineage tracing experiments are depicted in green and red to maintain the original mTmG reporter set up (i.e. a switch from tdTomato to eGFP).

### Software, statistics and online databases

The following software was used: Fiji (https://imagej.net/Fiji/Downloads) for image analysis (Schindelin et al., 2012), R (R Development Core Team, 2017) and R studio (https://rstudio.com/products/rstudio/download/) to generate the dotplots and violin plots in Figure 2 using the ggplot2 package (Wickham, 2009), ThermoFisher Cloud software and Microsoft Excel for qRT-PCR analysis using the ddCt method, LI-COR Image Studio Lite for Western blot analysis, Graphpad PRISM for generating graphs. Final figures were compiled in Adobe Illustrator. For Figure 1A, information was retrieved from http://www.findmice.org on the 7^th^ of January 2020.

---

**Supplementary Figure 1.**
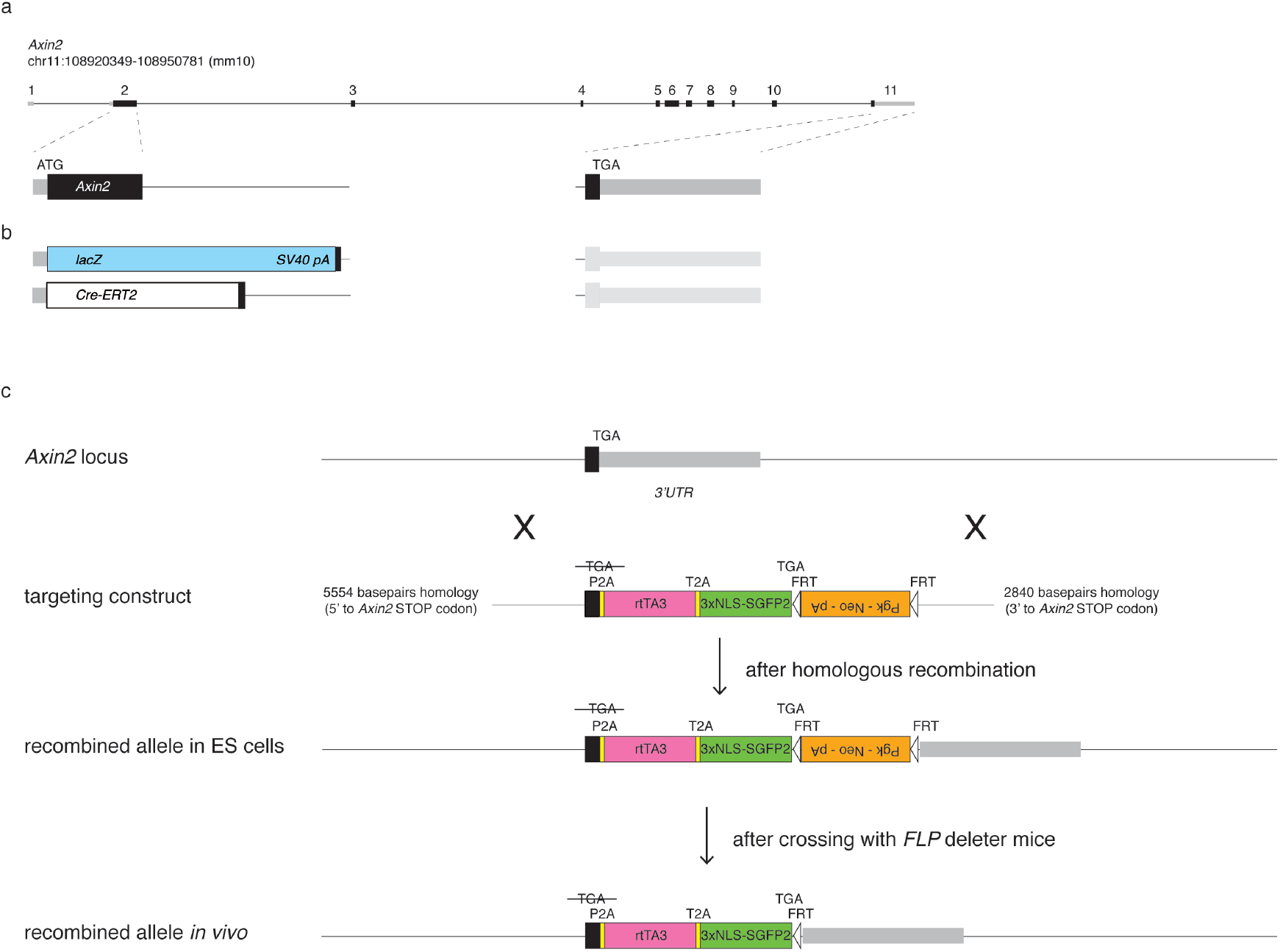
Details on the targeting strategy. (A) Schematic representation of the mouse *Axin2* locus (chr11:108920349-108950781, mm10 coordinates). (B) Most existing *Axin2* knock-in strains, including our previously generated *Axin2*^*CreERT2*^ driver and the frequently used *Axin2*^*lacZ*^ reporter, target the 5’ end of the gene by introducing the knock-in cassette at the start codon in exon 2. This disrupts the endogenous *Axin2* coding sequence. (C) Cartoon depicting details of the targeting construct and its homologous recombination into to the 3’ UTR of the *Axin2* locus. The targeting construct contained 5.6 kb (5’) and 2.8 kb (3’) homology arms. The *Axin2* stop codon (*TGA*) was mutated and immediately followed by the *P2A-rtTA3-T2A-3xNLS-SGFP2* multicistronic knock-in cassette. The *PGK-Neo-polyA* cassette, flanked by *FRT* sites, was cloned immediately downstream of the knock-in cassette in the reverse orientation and used for selection of embryonic stem cells carrying the recombined allele. After crossing founder mice with an *FLP* deleter strain, the recombined *in vivo* allele only carries a single remaining *FRT* site immediately downstream of the knock-in cassette, otherwise leaving the *Axin2* locus intact.

**Supplementary Figure 2.**
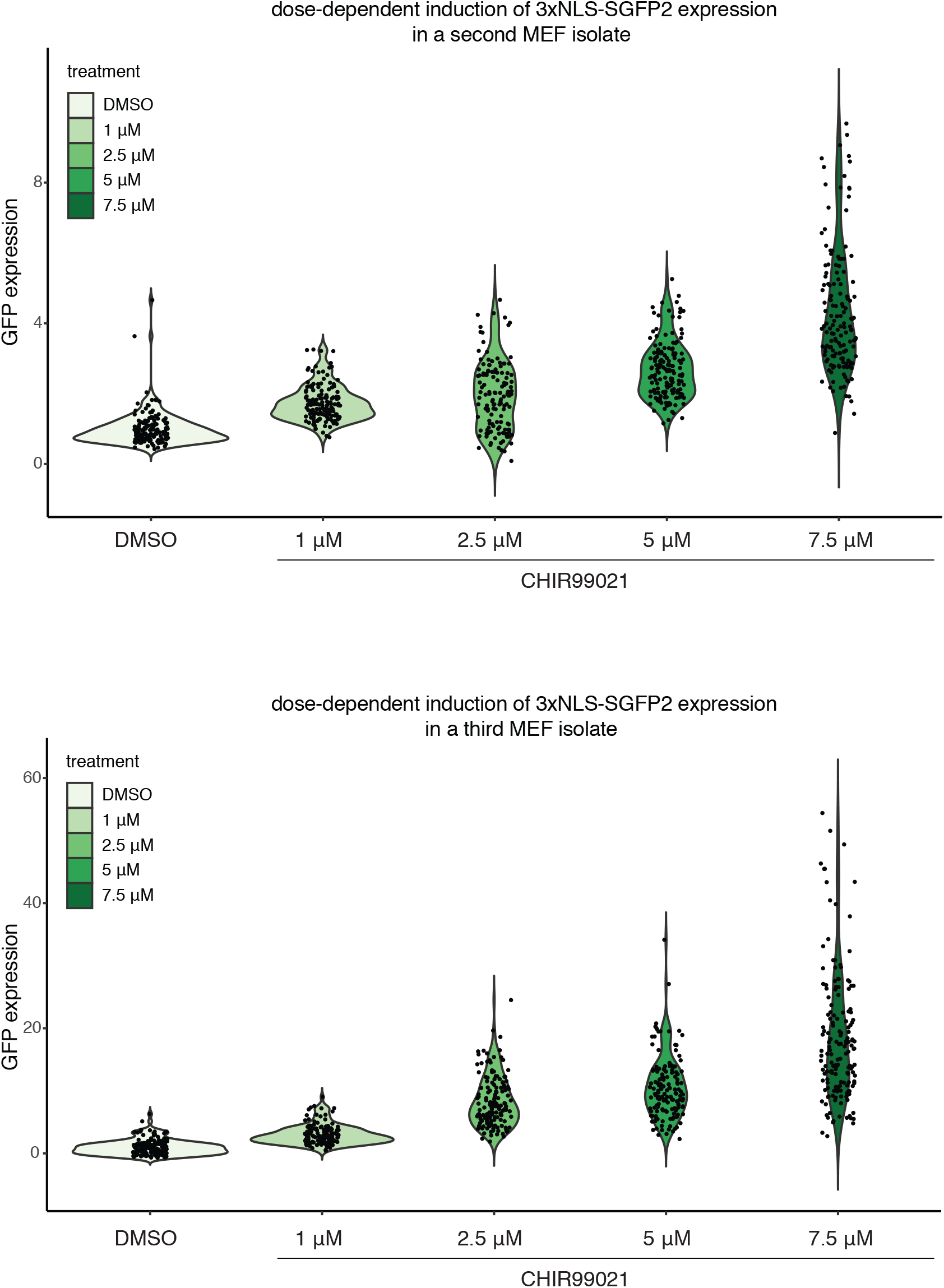
Dose-dependent induction of nuclear SGFP2 in MEFs. This figure shows the quantification of nuclear SGFP2 expression for two additional MEF isolates, supplementing the data shown in Figure 2G-H. Fixed MEFs were counterstained with DAPI to allow nuclear segmentation, after which the relative SGFP2 expression levels were calculated by correcting for the fluorescence intensity of the DAPI signal.

**Supplementary Figure 3.**
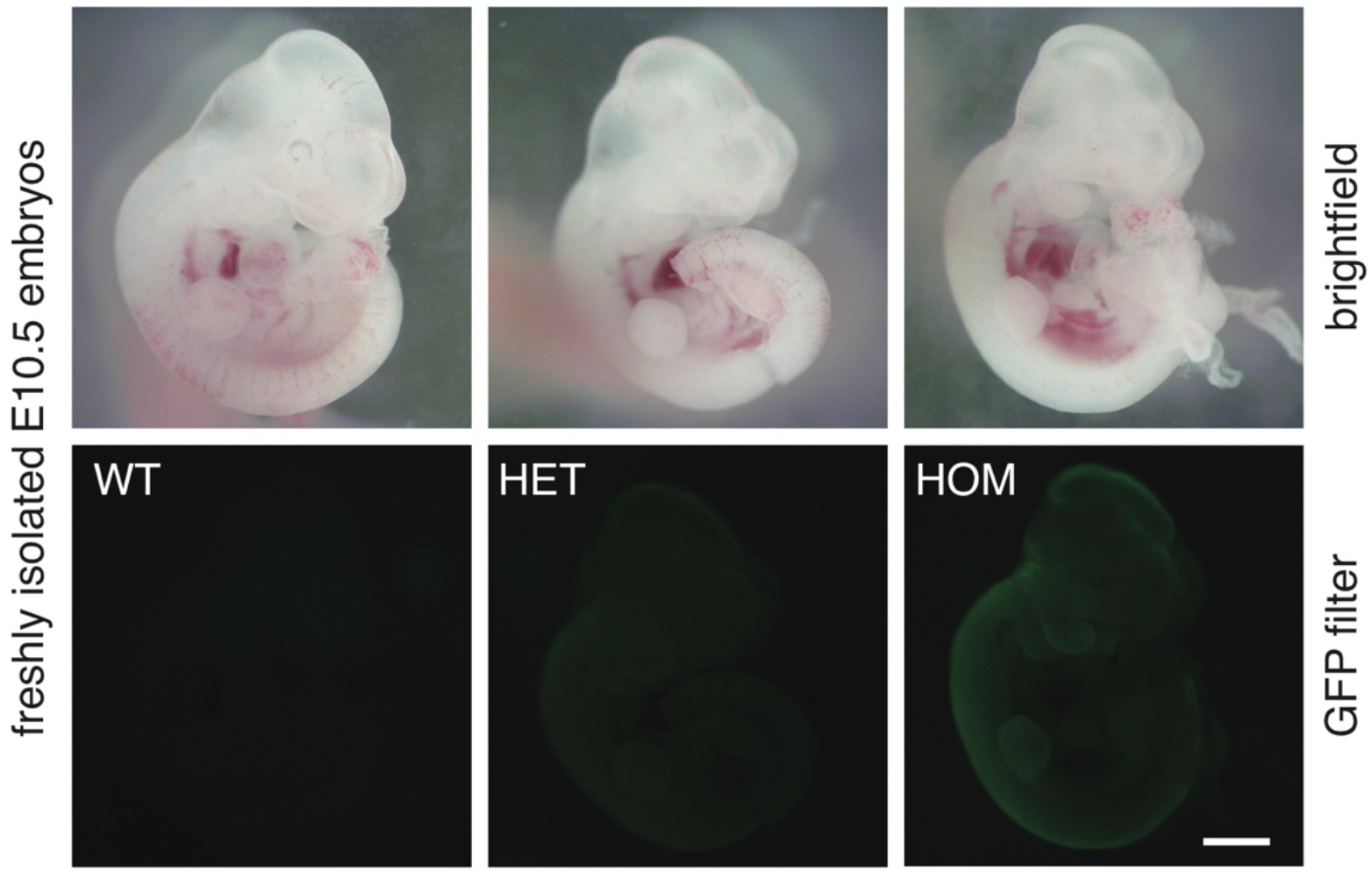
Wholemount detection of SGFP2 expression in *Axin2*^*P2A-rtTA3-T2A-3xNLS-SGFP2*^ embryos. Brightfield (top row) and fluorescent (bottom row) images of freshly isolated E10.5 wildtype (left), heterozygous (middle) and homozygous (right) embryos. A subtle difference in SGFP2 expression at the developing limb bud, the branchial arches, and along the dorsal neural tube, including the developing brain can be seen between heterozygous and homozygous embryos. Although we verified all genotypes by PCR, this difference in GFP intensity allowed us to distinguish wildtype, heterozygous and homozygous embryos by eye. All embryos depicted were imaged with the same exposure times (30 milliseconds for brightfield images, 8 seconds for a GFP long pass filter) under a dissecting microscope. Scalebar is 500 um. Note that size differences between embryos were not linked to the genotype, and thus most likely represent natural variation between individual embryos of the same litter.

**Supplementary Figure 4.**
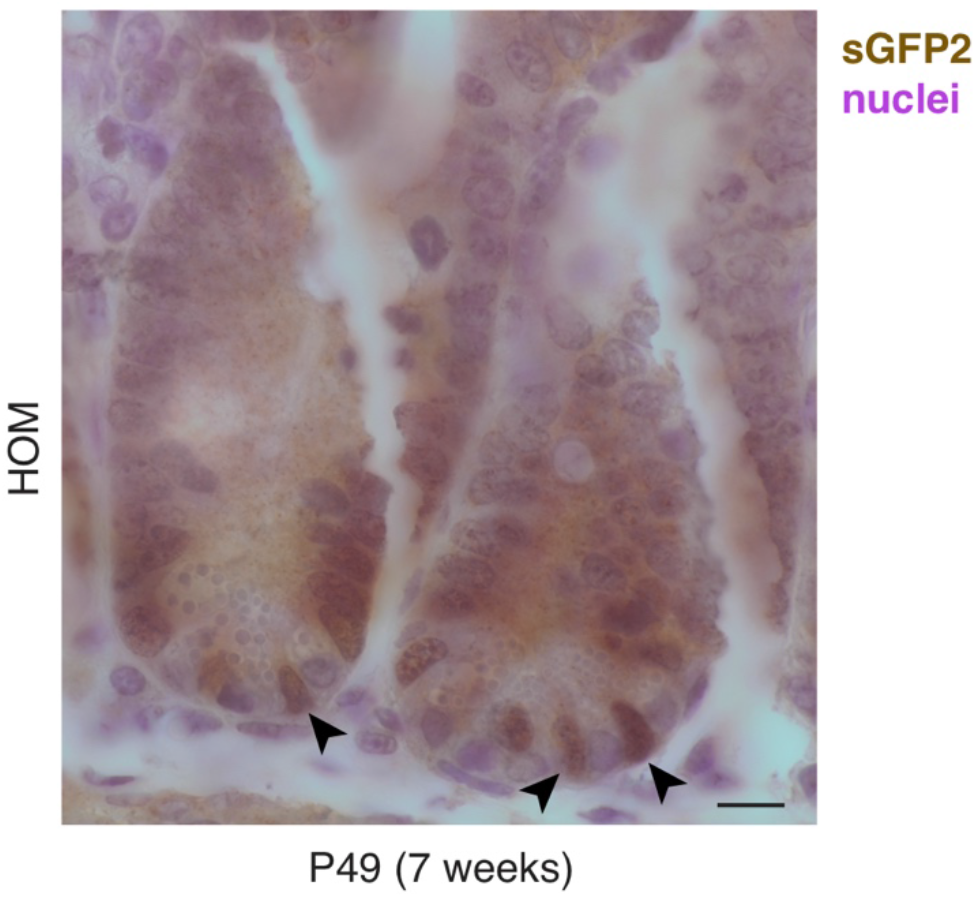
Immunohistochemical detection of SGFP2 in FFPE samples. Brightfield image showing the detection of nuclear SGFP2 in a formalin fixed, paraffin embedded (FFPE) tissue section from a homozygous *Axin2*^*P2A-rtTA3-T2A-3xNLS-SGFP2*^ adult (P49) animal. SGFP2 was visualized using an anti-GFP antibody and immunohistochemical detection. Nuclei are counterstained with hematoxylin. Scalebar is 10 um. Of note, we were not capable of robustly detecting SGFP2-positive cells postnatally in other organs tested (mammary gland, hair follicle and interfollicular epidermis of the skin and liver), which most likely reflects differences in the absolute levels of WNT/CTNNB1 signaling across tissues.

**Supplementary Figure 5:**
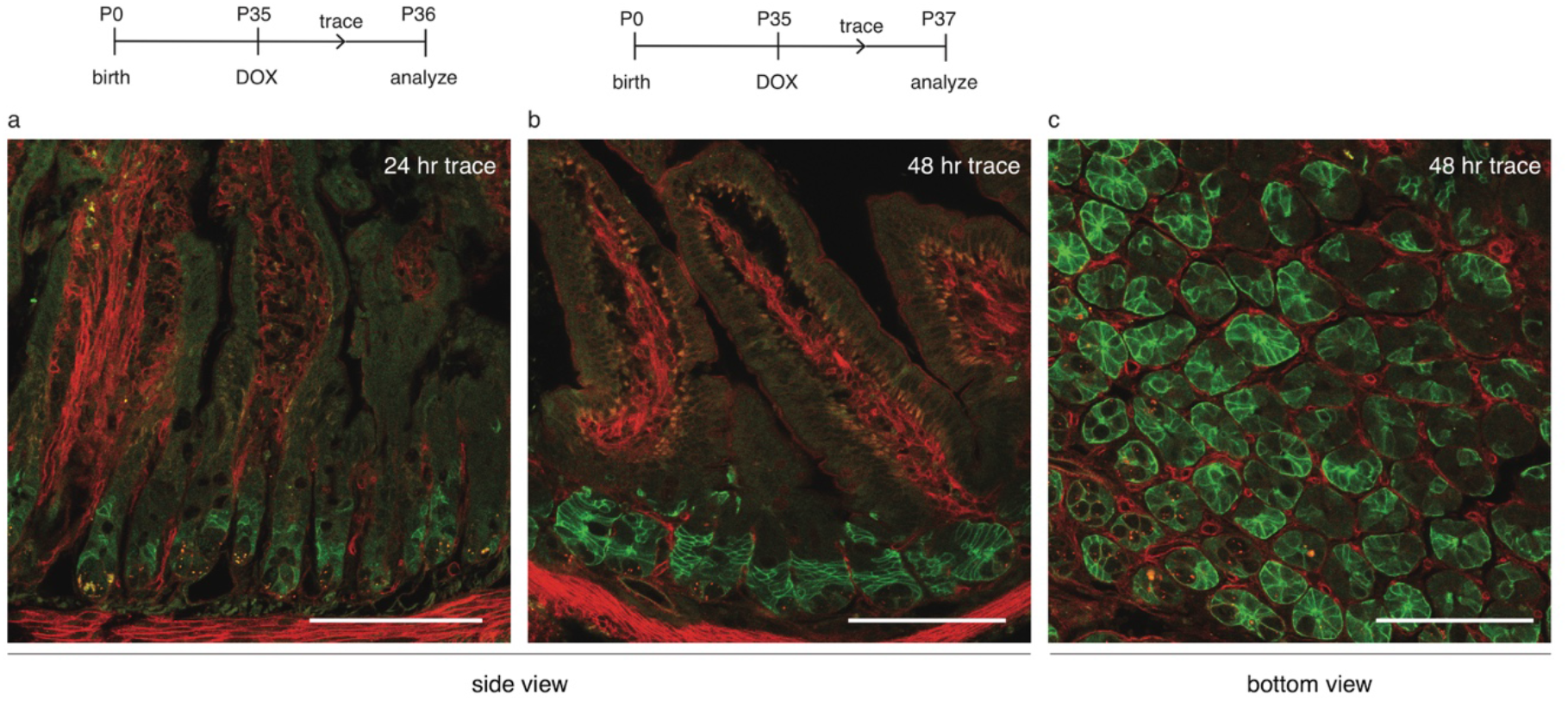
Short-term lineage trace of WNT/CTNNB1-responsive cells in intestinal tissue. Lineage tracing in WNT/CTNNB1-responsive cells was induced in pubertal mice at P35 via a single intraperitoneal injection of doxycycline (DOX) for 24 hours (A) or 48 hours (B-C). Experimental timelines are indicated on top. Wholemount confocal microscopy images showing a side view (A,B) and bottom view (C) of the intestinal epithelium from triple-heterozygous *Axin2*^*P2A-rtTA3-T2A-3xNLS-SGFP2*^*;tetO-Cre;Rosa26*^*mTmG*^ animals. The fluorescence signal is coming from the *Rosa26*^*mTmG*^ lineage tracing reporter allele, with the red color (mT) reflecting non-recombined cells and the green color (mG) reflecting recombined cells three days after induction. Recombined cells can be detected in the intestinal crypt as early as 24 hours after labeling in the intestinal (A). After 48 hours a substantial portion of cells in the crypt (B-C) can be detected with the concentration of DOX used. Scalebar is 100 *μ*m.

**Supplementary Figure 6:**
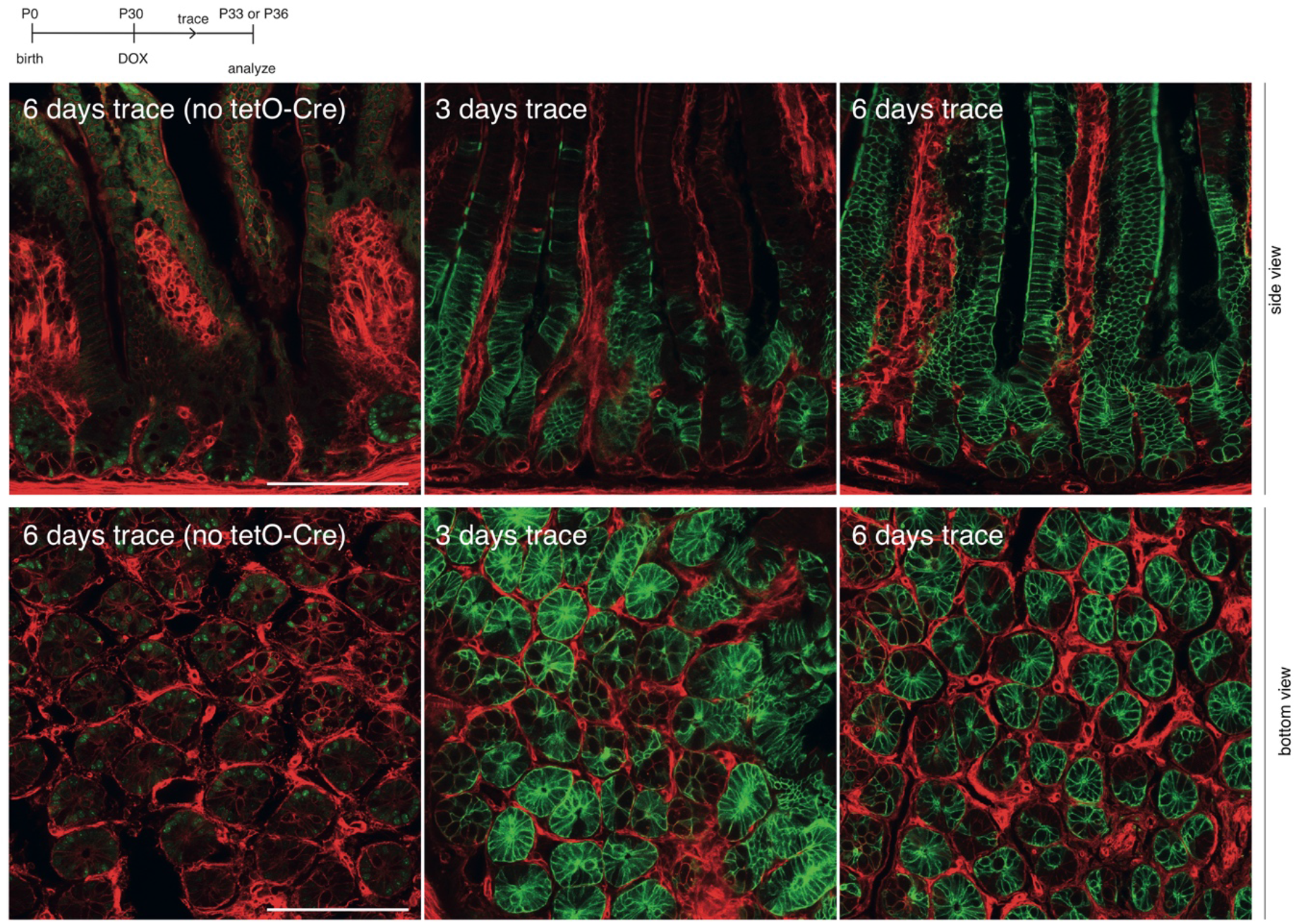
Lineage tracing of WNT/CTNNB1 responsive-cells in intestinal tissue. Lineage tracing in WNT/CTNNB1-responsive cells was induced in pubertal mice at P30 via a single intraperitoneal injection of doxycycline (DOX). A timeline for the experiment is depicted on top. Wholemount confocal microscopy images showing a side (top) and bottom view of the intestinal epithelium from triple-heterozygous *Axin2*^*P2A-rtTA3-T2A-3xNLS-SGFP2*^*;tetO-Cre;Rosa26*^*mTmG*^ animals. The WNT/CTNNB1-responsive lineage was traced for 3 and 6 days. The left panel shows a control sibling without TetO-Cre allele, in which only the 3xNLS-SGFP2 signal from the *Axin2* locus is visible. The middle panel shows the fluorescence signal coming from the *Rosa26*^*mTmG*^ lineage tracing reporter allele, with the red color (mT) reflecting non-recombined cells and the green color (mG) reflecting recombined cells 3 days after induction. The crypts show recombined cells, with the progeny of these stem cells starting to populate the villus compartment. The right panel shows recombined cells 6 days after induction of lineage tracing. Villi from recombined crypts are completely green. Scalebar is 100 *μ*m.

**Supplementary Figure 7:**
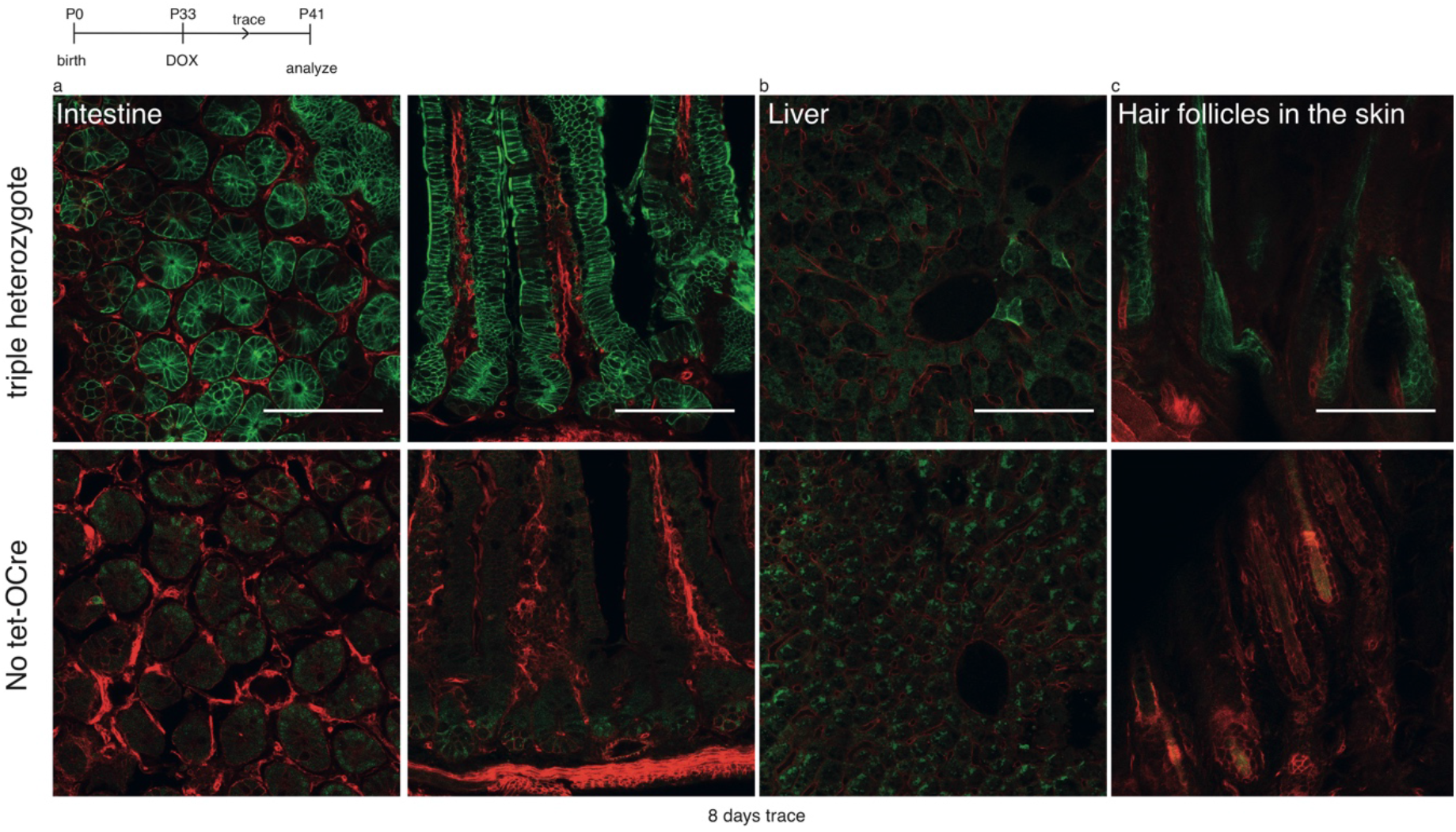
Lineage tracing of WNT/CTNNB1 responsive cell in postnatal tissues. Lineage tracing in WNT/CTNNB1-responsive cells was induced in pubertal mice at P33 via a single intraperitoneal injection of doxycycline (DOX). A timeline for the experiment depicted is shown on top. Wholemount confocal microscopy images showing different tissues from triple-heterozygous *Axin2*^*P2A-rtTA3-T2A-3xNLS-SGFP2*^*;tetO-Cre;Rosa26*^*mTmG*^ animals. The WNT/CTNNB1-responsive lineage was traced for 8 days. The bottom panel shows a control sibling without TetO-Cre allele where only the 3xNLS-SGFP2 signal from the *Axin2*^*P2A-rtTA3-T2A-3xNLS-SGFP2*^ is detectable. (A) Wholemount confocal microscopy images showing a bottom (left) and side (right) view of the intestinal epithelium. The top panels show the fluorescence signal coming from the *Rosa26*^*mTmG*^ lineage tracing reporter allele, with the red color (mT) reflecting non-recombined cells and the green color (mG) reflecting recombined cells. Both the crypt and villi compartments show a majority of recombined cells. (B) Wholemount confocal microscopy images of fixed and cleared liver, showing the recombined mTmG allele in sporadic cells adjacent to the central vein (top). (C) Wholemount confocal microscopy images of fixed and cleared skin, showing the recombined mTmG allele in multiple hair follicles (top). Scalebar is 100 *μ*m.

**Supplementary Figure 8.**
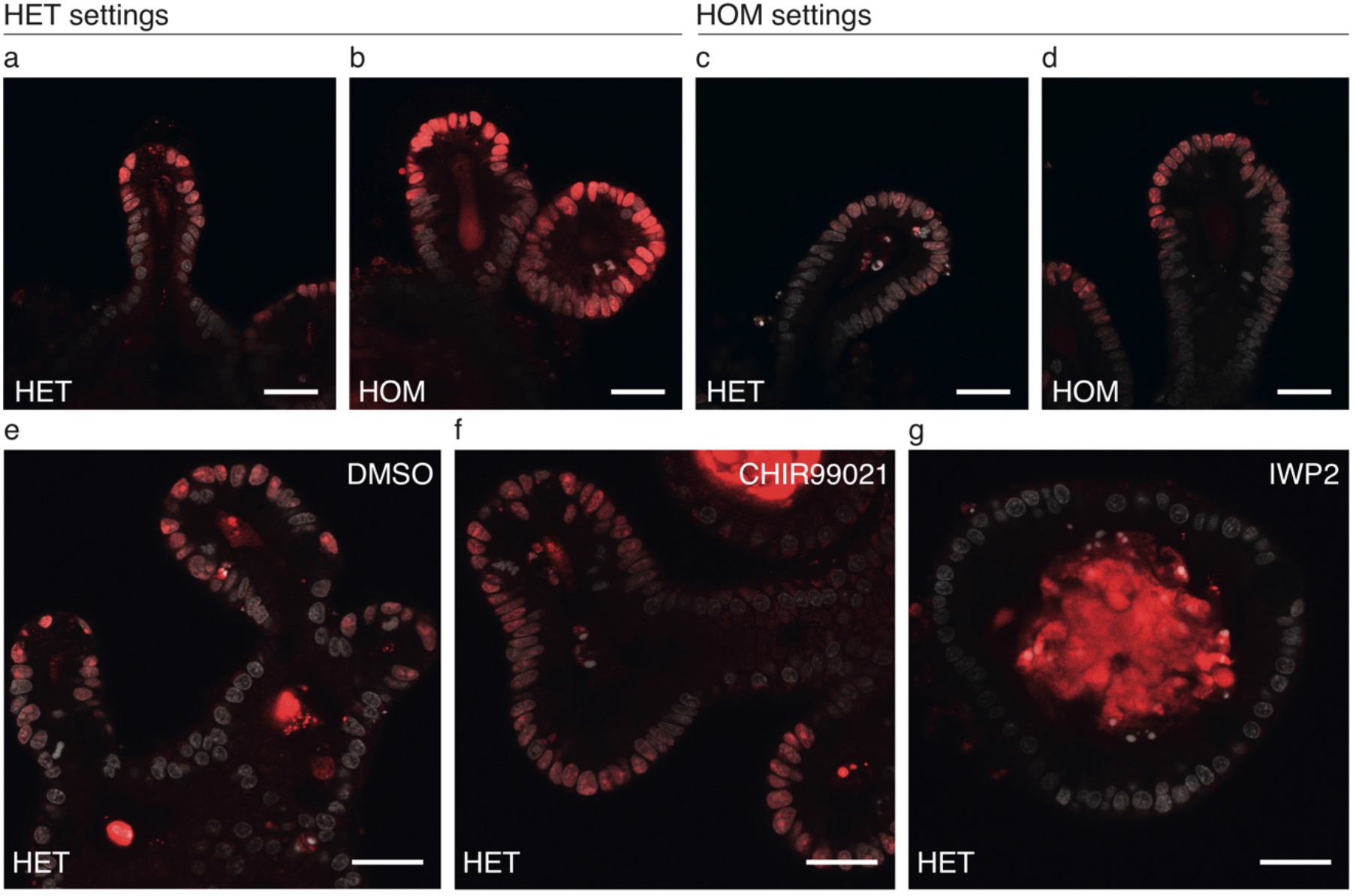
SGFP2 signal intensity in heterozygous versus homozygous small intestinal organoids. (A-D) Confocal microscopy images of crypts from fixed *Axin2*^*P2A-rtTA3-T2A-3xNLS-SGFP2 HOM*^ and *Axin2*^*P2A-rtTA3-T2A-3xNLS-SGFP2 HET*^ small intestinal organoids stained with nuclear dye DRAQ5. SGFP2 is depicted in red and DRAQ5 in grey. (A-B) were imaged with settings optimized for *Axin2*^*P2A-rtTA3-T2A-3xNLS-SGFP2 HET*^ organoids and (C-D) with settings optimal for *Axin2*^*P2A-rtTA3-T2A-3xNLS-SGFP2 HOM*^ organoids. Scalebar is 30 *μ*m. For panels (A-D) n=6 *Axin2*^*P2A-rtTA3-T2A-3xNLS-SGFP2 HET*^ organoids and n=7 *Axin2*^*P2A-rtTA3-T2A-3xNLS-SGFP2 HOM*^ were imaged in a single experiment. (E-G) Confocal microscopy images of crypts from fixed *Axin2*^*P2A-rtTA3-T2A-3xNLS-SGFP2 HET*^ small intestinal organoids stained with nuclear dye DRAQ5. Organoids were imaged with settings optimized for *Axin2*^*P2A-rtTA3-T2A-3xNLS-SGFP2 HET*^ organoids. Samples were treated with DMSO for 24 hours (E), CHIR99021 for 24 hours (F), or IWP2 for 48 hours (G) prior to fixation. Scalebar is 30 *μ*m. For panel (E-G) a total of 18 organoids were imaged in a single experiment (n=9 for DMSO, n=5 for CHIR99021, n=4 for IWP2). Representative images of all conditions are shown.

**Supplementary Movie 1. Nuclear SGFP2 expression in E12.5 heterozygous mammary bud**

3D rotation of an E12.5 mammary bud from a heterozygous *Axin2*^*P2A-rtTA3-T2A-3xNLS-SGFP2*^ embryo. Total Z-stack is 50 slices, corresponding to approximately 45 *μ*m depth.

**Supplementary Movie 2. Nuclear SGFP2 expression in E12.5 homozygous mammary bud**

3D rotation of an E12.5 mammary bud from a homozygous Axin2^P2A-rtTA3-T2A-3xNLS-SGFP2^ embryo. Total Z-stack is 44 slices, corresponding to approximately 30 *μ*m depth.

**Supplementary Movie 3. Traced *Axin2*-positive cells in the liver**

Wholemount confocal microscopy Z-stack through a piece of formalin fixed, cleared liver tissue from a triple-heterozygous *Axin2*^*P2A-rtTA3-T2A-3xNLS-SGFP2*^*;tetO-Cre;Rosa26*^*mTmG*^ mouse that was traced for 8 days after receiving a single intraperitoneal injection of doxycycline mid puberty. The fluorescence signal comes from the *Rosa26*^*mTmG*^ lineage tracing reporter allele, with the red color (mT) reflecting non-recombined cells and the membrane-localized green color (mG) reflecting cells that have recombined the reporter.

**Supplementary Table 1.**
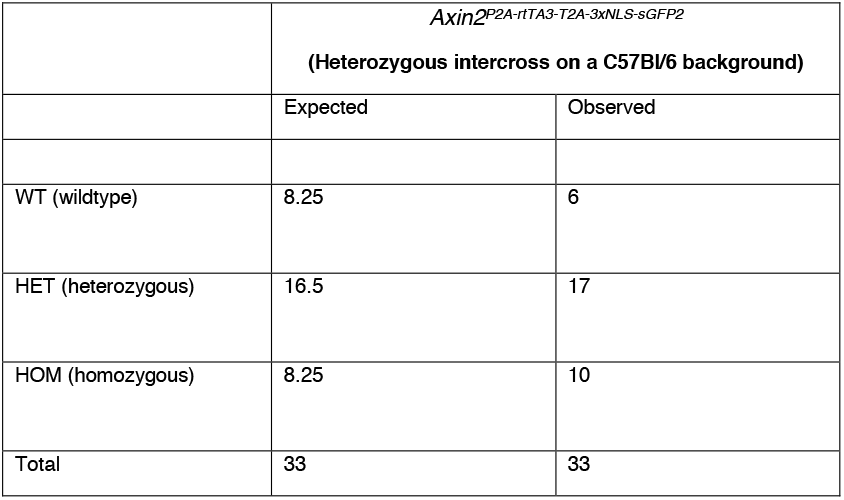
Mendelian inheritance of the 3’ *Axin2*^*P2A-rtTA3-T2A-3xNLS-SGFP2*^ allele. Homozygous *Axin2*^*P2A-rtTA3-T2A-3xNLS-SGFP2*^ mice are born at the expected mendelian ratios from heterozygous intercrosses on a C57BL/6 background. (Chi-square test, P=0.61 for both sexes combined).

**Supplementary Table 2.** Non-mendelian inheritance of the 5’ *Axin2*^*CreERT2*^ allele under some circumstances. Although we have bred *Axin2*^*CreERT2*^ mice without any trouble at multiple different animal facilities, we noticed that even heterozygous animals were born at sub-mendelian ratios after rederivation onto a C57BL/6J background. This phenotype was apparent during the first three generations of heterozygous backcrossing to FVB (Chi-square test, P=3.5×10^−6^ for both sexes combined), after which wildtype and heterozygous mice were again born at the expected ratios (Chi-square test, P=0.85 for both sexes combined).

|  | *Axin2*<sup>CreERT2</sup><br>(FVB backcross, generation F1-F3) |  |  | *Axin2*<sup>CreERT2</sup><br>(FVB backcross, generation F4-F6) |  |
| --- | --- | --- | --- | --- | --- |
|  | Expected | Observed |  | Expected | Observed |
| WT;WT (wildtype) | 41 | 62 |  | 14.5 | 14 |
| WT;KI (heterozygous) | 41 | 20 |  | 14.5 | 15 |
| Total | 82 | 82 |  | 29 | 29 |

**Supplementary Table 3.**
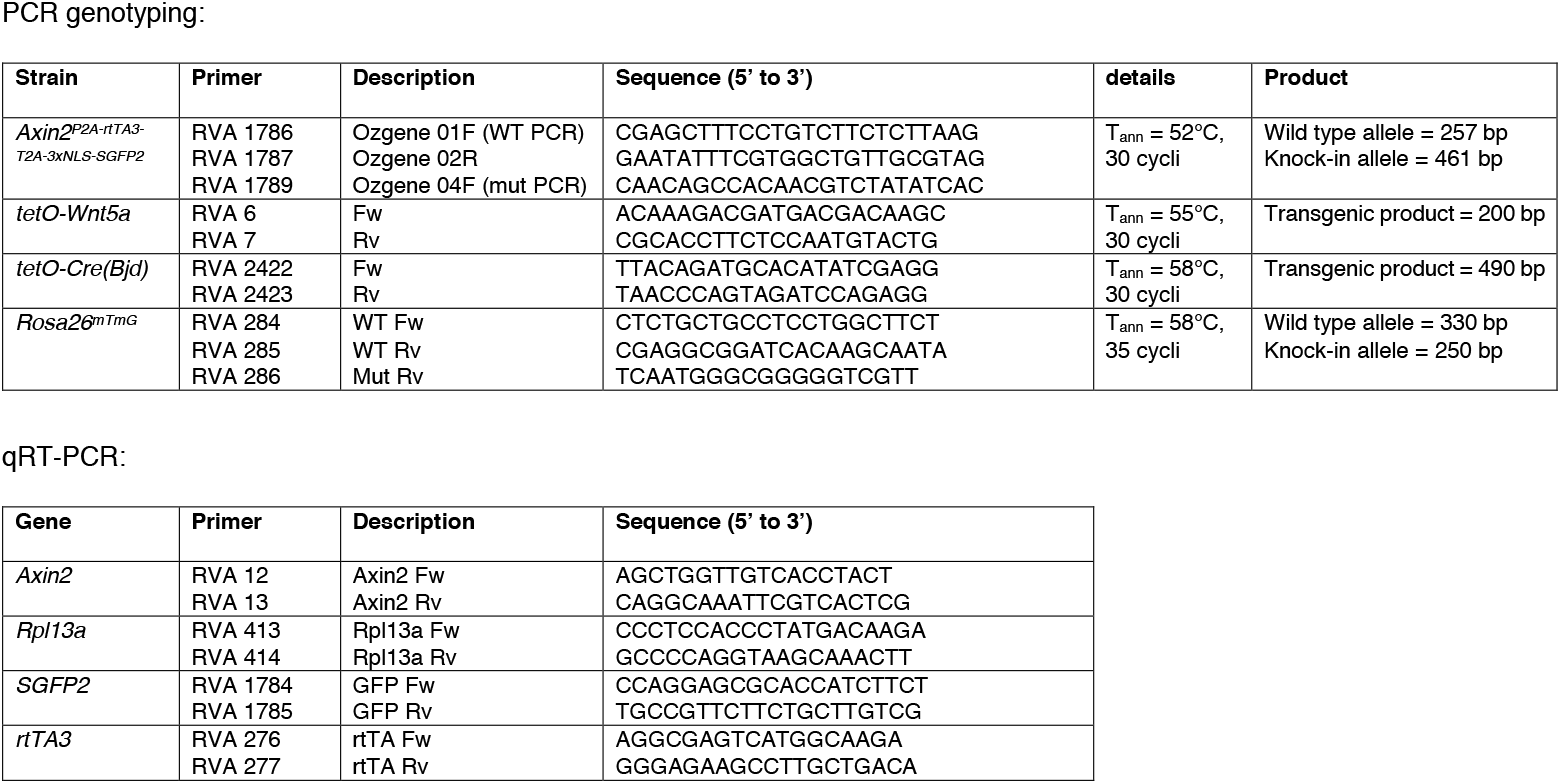
Primer sequences.

## Notes

http://shorturl.at/ilqIO

